# TSA-Seq Mapping of Nuclear Genome Organization

**DOI:** 10.1101/307892

**Authors:** Yu Chen, Yang Zhang, Yuchuan Wang, Liguo Zhang, Eva K. Brinkman, Stephen A. Adam, Robert Goldman, Bas van Steensel, Jian Ma, Andrew S. Belmont

**Affiliations:** Department of Cell and Developmental Biology, University of Illinois at Urbana-Champaign, Urbana, Illinois, 61801, USA.; Department of Bioengineering, University of Illinois at Urbana-Champaign, Urbana, Illinois, 61801, USA.; Computational Biology Department, School of Computer Science, Carnegie Mellon University, Pittsburgh, PA 15213; Division of Gene Regulation, Netherlands Cancer Institute, Plesmanlaan 121, 1016 HM Amsterdam, the Netherlands.; Department of Cell and Molecular Biology, Northwestern University Feinberg School of Medicine, Chicago, IL 60611; Center for Biophysics and Quantitative Biology, University of Illinois at Urbana-Champaign, Urbana, Illinois, 61801, USA.; Carl R. Woese Institute for Genomic Biology, University of Illinois at Urbana-Champaign, Urbana, Illinois, 61801, USA

**Author notes:** Corresponding Author: Andrew Belmont.

**Keywords:** epigenetics, epigenomics, genomics, nuclear structure, chromosome structure, nuclear speckles, LADs, transcription, nuclear compartments

## Abstract

While nuclear compartmentalization is an essential feature of three-dimensional genome organization, no genomic method exists for measuring chromosome distances to defined nuclear structures. Here we describe TSA-Seq, a new mapping method able to estimate mean chromosomal distances from nuclear speckles genome-wide and predict several Mbp chromosome trajectories between nuclear compartments without sophisticated computational modeling. Ensemble-averaged results reveal a clear nuclear lamina to speckle axis correlated with a striking spatial gradient in genome activity. This gradient represents a convolution of multiple, spatially separated nuclear domains, including two types of transcription “hot-zones”. Transcription hot-zones protruding furthest into the nuclear interior and positioning deterministically very close to nuclear speckles have higher numbers of total genes, the most highly expressed genes, house-keeping genes, genes with low transcriptional pausing, and super-enhancers. Our results demonstrate the capability of TSA-Seq for genome-wide mapping of nuclear structure and suggest a new model for nuclear spatial organization of transcription.

## Introduction

While the human genome has been sequenced, how this linear genome sequence folds in 3D within the nucleus remains largely unknown. New genomic methods such as Hi-C (Lieberman-Aiden et al., 2009; Rao et al., 2014) have generated increasing interest in how 3D chromosome folding may regulate genome functions during development or in health and disease. However, these 3C-based methods do not directly report on chromosome positioning within nuclei.

What is needed is an ability to translate microscopic views of DNA position relative to nuclear compartments (such as the nuclear lamina, nucleolus, or nuclear speckles) into genome-wide maps that show how close loci are to a given compartment and how the chromosomal fiber traverses between compartments. For example, whether transcriptionally active chromosome regions are targeted reproducibly to particular nuclear compartments has been a long-standing question. Using DNA FISH, a population-based, statistical shift towards the nuclear center has been observed for a number of genes undergoing transcriptional activation (reviewed in (Takizawa et al., 2008)), leading to the proposal of a gradient of increased transcriptional activity from the nuclear periphery to center (Bickmore, 2013; Takizawa et al. 2008). However, the functional significance of this radial positioning has been difficult to rationalize given the large variability of gene positioning within individual nuclei (Kölbl et al., 2012; Takizawa et al., 2008). Alternatively, this stochastic radial positioning of genes could be the consequence of a more deterministic positioning of genes relative to a nuclear compartment(s) that itself shows a stochastic radial positioning.

Nuclear speckles, excluded from the nuclear periphery and enriched towards the nuclear center (Carter et al., 1991), are an excellent candidate for such a nuclear compartment. Nuclear speckles were first visualized by transmission electron microscopy (TEM) as dense clusters of 20-25 nm diameter RNP (ribonucleoprotein) granules (Fakan and Puvion, 1980), termed “interchromatin granule clusters” (IGCs), and have alternatively been proposed to be storage sites for RNA processing components (Spector and Lamond, 2011) or transcription hubs for a subset of active genes (Hall et al., 2006; Shopland et al., 2003; Xing et al., 1995). Microscopic studies have demonstrated the very close association with (Moen, 2003; Xing et al., 1995) or even movement to (Hu et al., 2009; Khanna et al., 2014) nuclear speckles of a small number of genes upon transcriptional activation. One major problem, however, in judging the significance of this speckle-association has been the absence of any successful genome-wide survey of the prevalence of gene association with nuclear speckles. A low-throughput microscopy survey showed ~10/25 active genes with “close” association to nuclear speckles, but this small sampling of active genes might not be representative of the entire genome. Another significant problem has been the non-quantitative assessment of “close” used in previous studies and the absence of any comparison to the percentage of the genome localized within similar distances.

Current genomic methods such as DamID (Vogel et al., 2007) and ChIP-Seq (Landt et al., 2012) are limited in mapping nuclear speckle-associated domains as they measure molecular contact frequencies with particular proteins but not the actual cytological distances from specific nuclear compartments. Nuclear speckles behave like a dynamic phase-separated body (Brangwynne, 2011; Galganski et al., 2017; Zhu and Brangwynne, 2015) and no detectable DNA is contained within them (Spector and Lamond, 2011), while serial section TEM reconstructions showed large-scale chromatin fibers near, but not necessarily contacting, the periphery of these granule clusters (Belmont and Bruce, 1994). Thus the first challenge is detecting DNA sequences that are close to nuclear speckles but not in direct contact. Moreover, all known speckle proteins, although enriched in nuclear speckles, are also found at lower concentration throughout the nucleus; most of these proteins are involved in RNA processing and/or transcription and bound to chromatin or nascent transcripts. Therefore, the second challenge is producing a signal proportional to the total amount of a protein within a microscopic neighborhood rather than the small fraction of that protein in molecular contact with DNA.

Here we present “TSA-Seq”, the first genomic method capable of estimating cytological distances of chromosome loci genome-wide relative to a particular nuclear compartment and even inferring chromosome trajectories from one compartment to another. Our TSA-Seq results suggest a refined model of nuclear organization in which the proper metric is not fractional distance relative to the nuclear periphery versus nuclear center, but rather the distance of genomic regions to specific nuclear compartments- e.g. the nuclear lamina or speckles or still unknown regions, such as transcription factories.

## Results

### Gradient of diffusible free radical produced by TSA provides distance-dependent biotin labeling

To measure the relative distance of genomic regions to specific nuclear compartments, we turned to Tyramide Signal Amplification (TSA) (Wang et al., 1999), a commonly used immunochemistry method for amplifying signals for target detection. TSA uses an antibody-coupled horseradish peroxidase (HRP) to catalyze the formation of diffusible biotin-tyramide free radicals and create a free radical concentration gradient centered at the staining target (Figure 1A).

**Figure 1.**
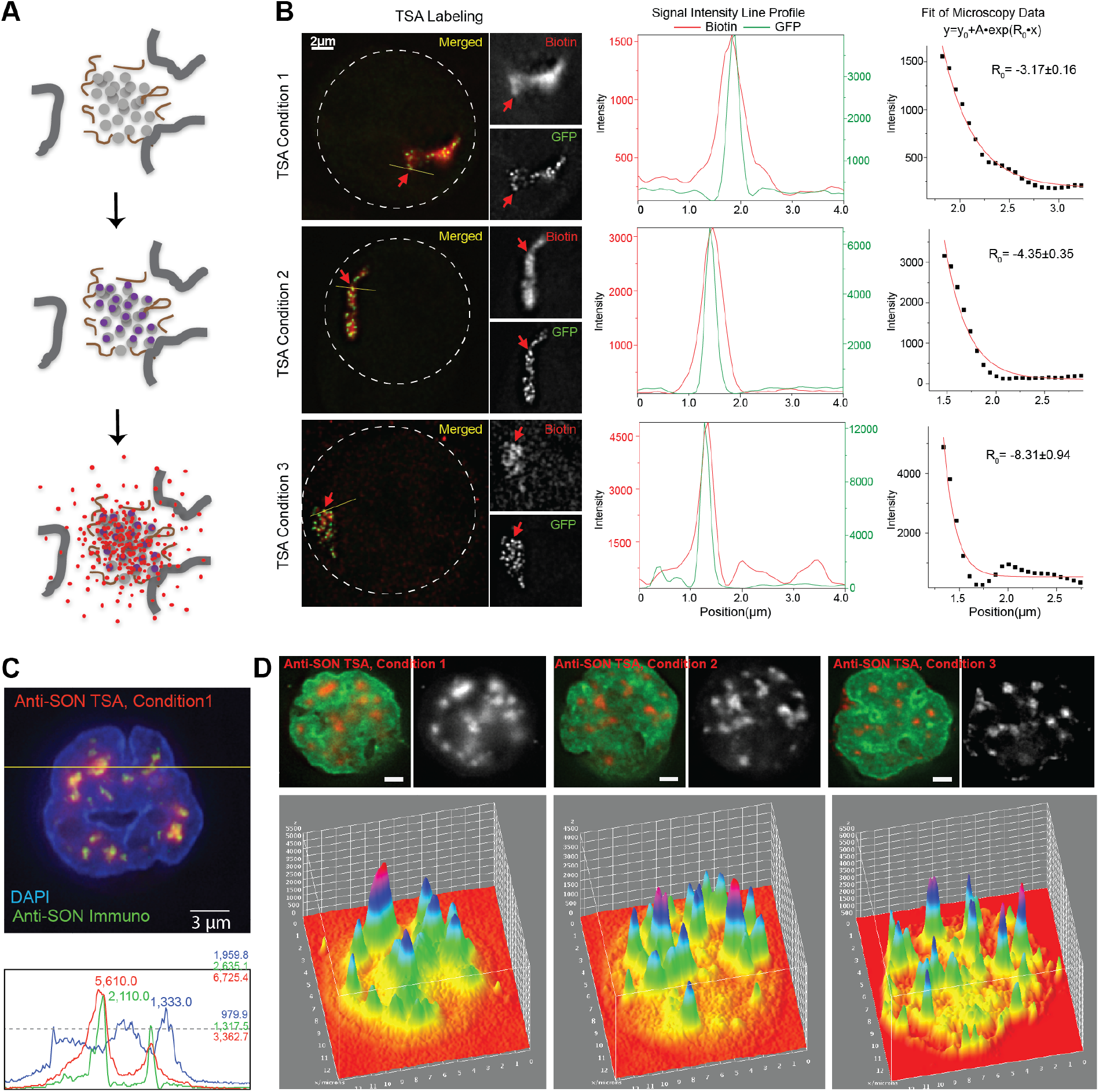
TSA generates biotin labeling which decays exponentially from the staining target. (A) Tyramide Signal Amplification (TSA): Primary and HRP-conjugated secondary antibodies (purple dots) target epitope (gray dots). HRP catalyzes formation of tyramide-biotin free radicals (red) which diffuse and covalently link to nearby proteins (gray dots), DNA (gray lines) and RNA (brown lines). (B) Exponential fitting of spreading of TSA staining: Condition 1 (top), Condition 2 (middle), and Condition 3 (bottom). Left: Biotin staining (red) of TSA staining of GFP-lac repressor (green) bound to BAC transgene array-white dashes mark nucleus. Middle: GFP spot (green) versus biotin intensities (red) along yellow line scan (left). Right: Exponential fit of biotin intensity versus distance from GFP spot. (C) Image (top) and intensities (bottom) along yellow line-scan (top) of: SON immunostaining (green), biotin staining after TSA (red), and DNA DAPI staining (blue). Scale bar: 3μm. (D) Images of SON TSA staining conditions I-III. Top: biotin staining (red) merged with DAPI DNA staining (green). Bottom: TSA staining surface plots. Scale bars: 2μm.

We measured how far the TSA signal spreads using a cell line with multiple intranuclear GFP spots produced by EGFP-lac repressor binding to 256mer lac operator DNA tags contained within an integrated BAC multi-copy transgene array (Hu et al., 2009). Each GFP spot served as a point source for biotin-tyramide free radical generation after anti-GFP TSA. Diffusion of tyramide free radicals from a point source combined with a spatially invariant free radical degradation rate is predicted to produce an exponential decay in free radical concentration with distance from this point source at steady-state (Wartlick et al., 2009). Using the line profiles of streptavidin staining of the observed biotin distribution, we fit the observed signals to exponential decay functions (Figure 1B). The unmodified TSA reaction product spread over an ~ 1 μm radius (Condition 1). Adding sucrose to increase the solution viscosity reduced this spreading moderately (Condition 2), although it reduced the biotin labeling ~4-fold (data not shown). Adding both sucrose and DTT (Condition 3), to reduce the tyramide free radical’s lifetime and diffusion path length, reduced biotin labeling ~20-fold (data not shown) but reduced the staining radius less than ~0.5 μm.

To label nuclear speckles, we used anti-SON (Ab1, Methods) antibody staining against the protein SON, a highly specific marker for nuclear speckles (Figure S1A). Comparison of anti-SON immunostaining (Figure 1C, green) with the streptavidin labeling of the TSA tyramide-biotin signal (Figure 1C, red), shows TSA signal spreading over distances consistent with our estimates from the GFP-stained lac operator arrays (Figure 1B). The intranuclear biotin labeling is dominated by this TSA signal spreading from nuclear speckles, as shown by 3-dimensional surface plots (Figure 1D) for Conditions 1-3, consistent with the low nucleoplasmic SON immunostaining (Figures 1C and S1A). TSA reactions using no primary antibody showed minimal labeling (data not shown).

### TSA-Seq translates the distance-dependent, TSA biotin-labeling into a quantitative sequencing readout

To convert TSA staining into a genomic readout, we followed the TSA reaction with reversal of formaldehyde cross-linking, DNA isolation, pull-down of biotinylated DNA, and high-throughput sequencing (Figure 2A). Workflow quality control included microcopy of strepavidin-stained cell aliquots following the TSA reaction and anti-biotin dot-blot staining of the purified DNA (data not shown). Previous TSA applications, including an attempt to identify DNA sequences localizing near PML bodies (Ching et al., 2013), have focused on protein labeling, through the known reaction of tyramide free radicals with tyrosines and tryptophans. We found that TSA also labels DNA (data not shown). Using DNA labeling we avoid potential biases introduced by differential chromatin solubilization and/or variations in the tyrosine and tryptophan content of surrounding non-histone proteins. As compared to ChIP-Seq, input variation across the genome is minimized and pull down specificity can be improved using harsher elution conditions.

**Figure 2.**
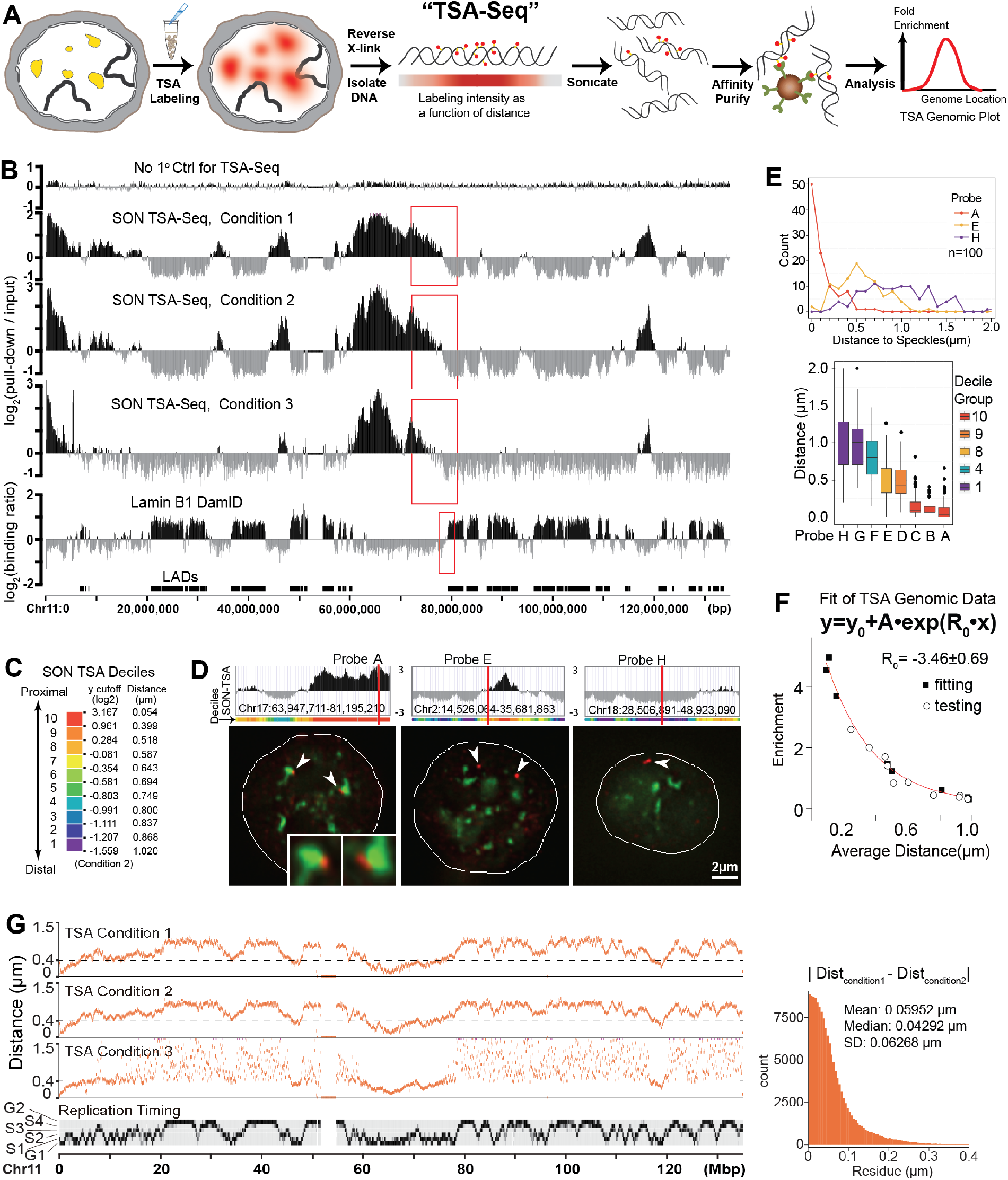
Genome-wide mapping of cytological distance from nuclear speckles using TSASeq. (A) Workflow schematic; (B) SON TSA-Seq genomic plots (human Chr11): top to bottom- no primary control, TSA staining Conditions 1-3, lamin B1 DamID. Red boxes-SON TSA-Seq peak-to-valley versus inter-LAD -LAD transitions. (C) Color-coding, TSA-Seq scores and estimated speckle distances for 10 SON TSA-Seq deciles (TSA condition 2). (D) Top: SON TSA-Seq scores and probe locations for peak (probe A), transition (probe E), and valley (probe E) regions. Bottom: 3D immuno-FISH-SON (green), FISH (red), nucleus (white lines). Scale bar: 2μm. (E) Histograms (top) and box plot (bottom) of probe distances to nearest speckle (n=100). (F) Exponential fit (red line) of SON TSA-Seq enrichment relative to mean (Condition 1) versus speckle distance calculated from 8 probes (black squares); 10 additional probes (open circles) map close to fitted curve. (G) Left (top to bottom): speckle distances (chr11) calculated from inverse of exponential fit (Conditions 1-3) versus Repli-Seq DNA replication timing. Right: histogram of distance residuals between Conditions 1 and 2.

We generated TSA enrichment genomic maps by plotting the log_2_ ratio of the normalized pull-down read density versus the normalized read density of input DNA over a 20 kbp sliding-window. We observed very similar enrichment maps for all three staining conditions (Figure 2B), with an ~16-32-fold enrichment/depletion dynamic range. Enrichment peaks were slightly sharper and the dynamic range higher for staining Condition 2, which produced less spreading of the tyramide free radical. Condition 3 further sharpened larger TSA-Seq peaks but reduced or eliminated smaller peaks and flattened many valleys, as expected by the more limited range of the free radical diffusion. The no primary control TSA labeling produced a nearly flat map.

Valleys in the SON TSA-Seq map align with lamina-associated domains (LADs) identified by lamin B1 DamID (Figure 2B), consistent with the known depletion of nuclear speckles near the nuclear periphery (Carter et al., 1991). Unlike the relatively abrupt transitions between inter-LADs and LADs, defined by the DamID molecular proximity mapping method, SON TSA-Seq maps typically show gradual, approximately linearly decreasing signals over the several Mbps separating SON TSA-Seq peaks and valleys which then levels close to the corresponding lamin B1 inter-LAD to LAD boundary (Figure 2B, red rectangles). We obtained nearly identical TSA-Seq maps using a different anti-SON antibody (Ab2, Methods) as well as a monoclonal antibody to a different speckle marker (phosphorylated SC35) (Figure S1B&C). These speckle TSA-Seq maps are distinctly different from published SC35 and SON ChIP-Seq results, showing near uniform levels at Mbp-scale resolution and localized peaks over cis-regulatory regions at the resolution level of individual genes (Figure S1D) (Ji et al., 2013; Kim et al., 2016). In contrast to TSA-Seq, which measures cytological distance to the staining target regardless of whether the proteins are within crosslinking distance, ChIP-Seq as a molecular proximity method detects only the small fraction of SC35 or SON likely associated over nascent transcripts within cross-linking distance to DNA. Indeed, these published ChIP-Seq data sets show localized peaks over all active genes, regardless of their distance to nuclear speckles (Figure S1E).

### SON TSA-Seq score is proportional to cytological distance from nuclear speckles and can be calibrated to estimate mean distance in μm

We used 3D immuno-FISH to validate that SON TSA-Seq signals are proportional to distance from nuclear speckles. We measured the distances to the nearest speckle using 8 BAC FISH probes corresponding to three peaks, three valleys, and two transition regions, as visualized in SON TSA-Seq color-coded decile and genomic plots (Figures 2C, 2D and S2A-S2C, Probes A-H). A strikingly deterministic localization near nuclear speckles was seen for probes A-C, corresponding to large SON TSA-Seq peaks. Nearly 100% of alleles probed mapped within 0.5 μm of a speckle, with mean distances between ~0.1-0.2 μm. Probes from transition (D-E) and valley (F-H) regions showed progressively larger mean speckle distances and broader distance distributions with decreasing SON TSA-Seq signals (Figures 2E and S2A-S2C).

If the SON TSA signal is dominated by spreading of tyramide free-radicals from nearby speckles, then the TSA-Seq dependence of biotinylated DNA enrichment as a function of speckle distance should follow a similar exponential fit as observed by microscopy of TSA staining (Figure 1B). Indeed, we can closely fit an exponential curve (Figures 2F and S2D, red lines) to the plot of DNA genomic enrichment versus mean speckle distance for DNA probes A-H (black squares). For each TSA staining condition, the estimated decay constants from genomic enrichment plots matched the decay constant calculated from the microscopy of TSA-labeled lac operator sites within the estimated fitting error (Figure S2E). To test how well this exponential fit would apply to other genomic regions, we acquired new FISH speckle distance measurements for an additional 10 BAC FISH probes targeting different genomic regions (Figure S2, Probes IR). Plotting mean speckle distance versus TSA genomic enrichment for these additional FISH probe regions showed a close match to the original fitted curve (Figure 2F, open circles, 0.039 (0.035) μm mean (median) difference between measured versus predicted distances).

To further test how well TSA-Seq could estimate mean distance genome-wide, we used these exponential fits to convert TSA enrichment to an estimated mean distance. We observe a very high reproducibility across the entire genome comparing calculated distances from Condition 1 versus Condition 2 SON TSA-Seq signals (residual mean, median, sd: 0.060, 0.043, 0.063 μm, Figure 2G). Reproducibility is also high for Condition 3 for predicted mean speckle distances less than ~0.4 μm (Figure 2G, left panel). Above 0.4 μm, speckle distance estimation is noisy because the SON TSA genomic enrichment decay curve has reached a plateau equal to background (Figure S2D), which also explains the flattening of valleys observed in Condition 3 SON TSA-Seq genomic plots (Figure 2B).

### Applying TSA-Seq to other nuclear compartment markers provides integrated view of nuclear organization that mirrors microscopic view

Our quantitative model for how TSA-Seq works predicts that any speckle marker with similar immunofluorescence speckle staining as SON, regardless of whether they are colocalized at a molecular level, should show a very similar TSA-Seq pattern. Indeed, the signals for the SC35 and SON TSA-Seq are nearly identical (Figures S1C, 3A and 3B).

**Figure 3.**
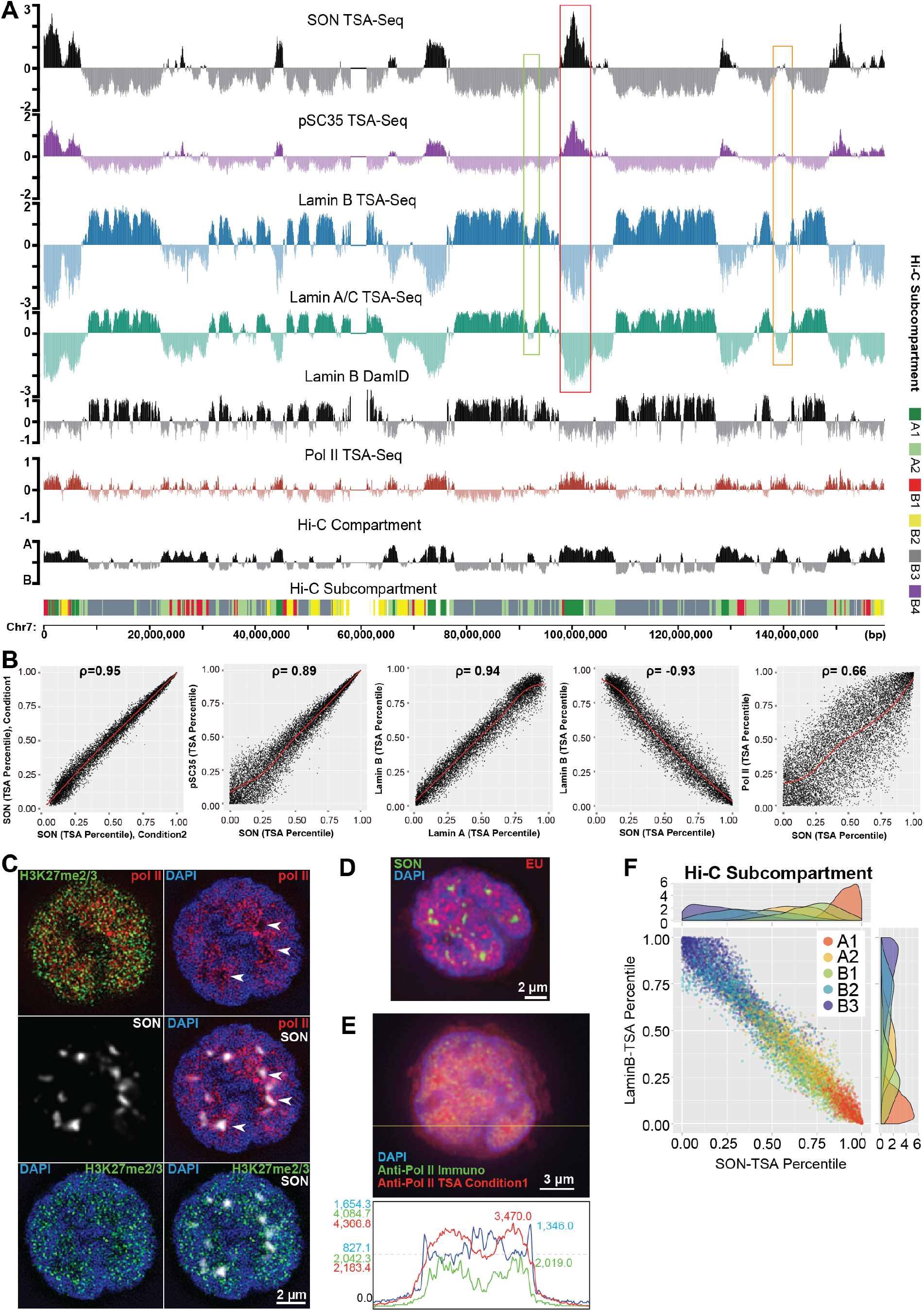
TSA-Seq versus light microscopy view of nuclear organization. (A) (top to bottom) TSA-Seq (chr7) for SON, phosphorylated SC35, lamin B, lamin A/C, RNA Pol II versus lamin B1 DamID, Hi-C compartment eigenvector and subcompartment assignment. Speckle TSA-Seq large peaks (red boxes), small peaks (orange boxes), and “peaks-within-valleys” (green boxes) align to lamin TSA-Seq deep valleys, shallow valleys, and dips between peaks, respectively. (B) 2D TSA-Seq percentile scatter-plots for (left to right): same target SON but two TSA labeling conditions, SON versus pSC35, lamin A/C versus lamin B, lamin B versus SON, RNA Pol II versus SON TSA (300 kb bins). (C) Top: 3D SIM of RNA pol II (red) versus H3K27me2/3 (left), RNA pol II (red) relative to DAPI (blue) (middle), and H3K27me3 (red) relative to DAPI (blue) (right). Bottom: SON wide-field image (left) superimposed on 3D SIM images (middle, right). Scale bar: 2 μm. (D) EdU 5 min pulse-label of nascent RNA (red) versus DNA (DAPI) staining and SON immunofluorescence (green). Scale bar = 2 μm. (E) Image (top) and aligned intensities (bottom) along line-scan (yellow, top) of RNA pol II immunostaining (green) versus TSA staining (red) and DAPI (blue). RNA pol II TSA signal (red) is high over nuclear interior but lower near nuclear periphery (and nucleoli). Scale bar: 3μm. (F) 2D lamin B-SON TSA-Seq scatter plots showing distribution of Hi-C subcompartments (320 kb bin size).

To further test the potential for TSA-Seq to probe nuclear organization, we performed TSA-Seq against additional nuclear compartment markers. We chose lamin A and lamin B (LB1 and LB2) as targets to further compare the relationship between chromosome distances to the nuclear lamina versus speckles. We also chose RNA pol II as a general marker for active nuclear domains to compare with the speckle TSA-Seq chromosome patterns.

Lamin A and B TSA-Seq are very similar and strikingly inversely correlated with speckle TSA-Seq throughout most of the chromosome regions-peaks, valleys, and the linear trajectories connecting peaks with valleys (Figures 3A and S3A). Whereas the lamin B1 and DamID signal shows a more binary distribution, lamin TSA-Seq shows distinctly different valley depths for different inter-LADs regions. Deeper valleys are correlated with larger SON TSA-Seq peaks (Figures 3A and S3B, boxes). The genome-wide inverse correlation between lamin and speckle TSA-Seq percentiles, defining a “nuclear lamina-speckle axis”, is demonstrated by the 2D scatterplot of lamin B versus SON TSA-Seq (Figure 3B).

RNA pol II TSA-Seq shows a lower dynamic range and more binary signal compared to SON TSA-Seq (Figure 3A). In contrast to SON TSA-Seq triangular-shaped peaks of varying heights, RNA pol II TSA-Seq shows broader, plateau-shaped peaks of similar heights. The correlation between SON and RNA pol II TSA-Seq is considerably lower than the correlations seen between related data sets such as lamin A versus lamin B TSA-Seq or SON versus SC35 TSA-Seq (Figure 3B). Instead, the RNA pol II TSA-Seq chromosomal profile strikingly matches the Hi-C eigenvector (Figure 3A), delineating the A (positive) and B (negative) Hi-C compartments known to be correlated with inter-LADs and LADs, respectively.

Comparing the TSA-Seq browser views with direct microscopy visualization of DNA, RNA pol II, SON, and H3K27me3 (Figure 3C) provides an integrated view of nuclear organization.

RNA pol II and nascent transcript foci are largely excluded from a thin zone at the nuclear periphery, but are distributed throughout most of the nuclear interior, although they surround and may even be concentrated near nuclear speckles (Figure 3C&D). The close spacing of these foci and spreading of the TSA staining produces a merged, low contrast RNA pol II TSA image. This relatively uniform RNA pol II TSA staining pattern (Figure 3E) - higher intensity in the nuclear interior and lower intensity in a peripheral zone close to the nuclear periphery (Figure 3E) - explains the low and flat RNA pol II TSA-Seq peaks and valleys overlapping with inter-LADs and LADs, respectively (Figure 3A).

At higher resolution, large SON TSA-Seq peaks align only with a subset of RNA pol II TSA-Seq positive regions, consistent with the RNA pol II immunostaining result: although there is a high concentration of RNA pol II that forms “rings” around the speckle periphery (Figure 3C, white arrow heads), there are also many RNA pol II foci widely distributed throughout the nuclear interior which are not speckle associated. Larger SON TSA-Seq speckle peaks align with deeper lamin A and B TSA-Seq valleys, consistent with the more interior localization of nuclear speckles. Most of the smaller SON TSA-Seq local maxima, including small peaks and “peaks within valleys”, align with the remaining RNA pol II TSA-Seq positive regions. These smaller SON TSA-Seq local maxima correlate with small dips or shallow valleys in the lamin TSA-Seq. Thus they are predicted to protrude smaller distances into the nuclear interior, consistent with the distribution of RNA pol II foci within several hundred nm of the nuclear periphery with which these active chromosomes regions may be interacting. Interestingly, the RNA pol II positive regions that contain large SON TSA peaks overlap extensively with A1 Hi-C subcompartment regions defined in GM12878 cells, which have a similar Hi-C profile to K562 cells (Rao et al., 2014). In contrast, the remaining RNA pol II positive regions, which contain smaller SON TSA-Seq local maxima, are contained within A2 Hi-C subcompartment regions.

Positive lamin B1 DamID regions (LADs) and lamin TSA-Seq peaks overlap with B2 and B3 repressive Hi-C subcompartments. In contrast, polycomb-associated, repressive B1 Hi-C subcompartments are interspersed between the two types of active Hi-C subcompartments and to a lesser extent between B2 and B3. This B1 subcompartment genomic distribution is consistent with the microscopic interspersed foci of RNA pol II and H3K27me3, a marker of polycomb-repressed chromatin, throughout the nuclear interior and the closer approach of H3K27me3 foci to the nuclear periphery (Figure 3C).

A genome-wide view of the relative nuclear spatial distributions of these different Hi-C subcompartments along a nuclear lamina-speckle axis is provided by 2D scatterplots in which each dot, representing a 300 kb chromosome segment, is plotted as a function of lamin B and SON TSA-Seq percentiles (Figure 3F). Interestingly, we found a more peripheral distribution of B3 versus B2 sub-compartments, hinting at a heterogeneity within or between LADs.

### TSA-Seq predicts intra-nuclear chromosome trajectories, linking genomic analysis with microscopy investigations of nuclear organization

Microscopy-based investigations of intra-nuclear chromosome topology have previously been limited by their low-throughput. Here we show the ability of TSA-Seq to predict chromosome trajectories from genome-wide data, and thus guide microscopy-based examination of chromosome organization.

The several Mbp genomic distances between SON TSA-Seq valleys and peaks (Figure 2B, red boxes) are consistent with the micron-scale distances of nuclear speckles from the nuclear periphery and the previously observed ~1-3 Mbp per micron compaction of large-scale chromatin fibers (Hu et al., 2009). We predict that the linear transitions between neighboring peaks and valleys seen in the SON TSA-Seq log2 scores correspond to linear chromosome trajectories that stretch from the nuclear periphery to a nuclear speckle.

To test that the TSA-Seq map can indeed predict both chromosome location and chromosome topology, we first selected a 4 Mbp transition from SON TSA-Seq valley to peak for validation by immuno-FISH. Using two probes located near the LAD boundary (#1) and mid-transition (#3), we observed 77% of alleles showing probe #1 (red) near the nuclear periphery and probe #2 (yellow) towards the interior, with 96% of examples showing an obvious peripheral to interior orientation for probes #1 versus 3 (Figure S4A). Using 4 probes (red), spanning from the LAD boundary to within 1 Mbp of the speckle peak, we observed curvilinear trajectories typically beginning near the periphery and pointing towards a nuclear speckle (green) (Figure S4B). Adding a 5^th^ probe (yellow) located at the SON TSA-Seq peak, showed that this trajectory terminates at the speckle periphery (Figure 4A). Similar trajectories were observed for a second 3 Mbp valley-to-peak transition (Figure 4B). A probe set covering a complete 11-12 Mbp valley-to-peak-to-valley SON TSA-Seq signal revealed “U” or “V” chromosome trajectories from the nuclear periphery to a speckle and back to the nuclear periphery (Figure 4C).

**Figure 4.**
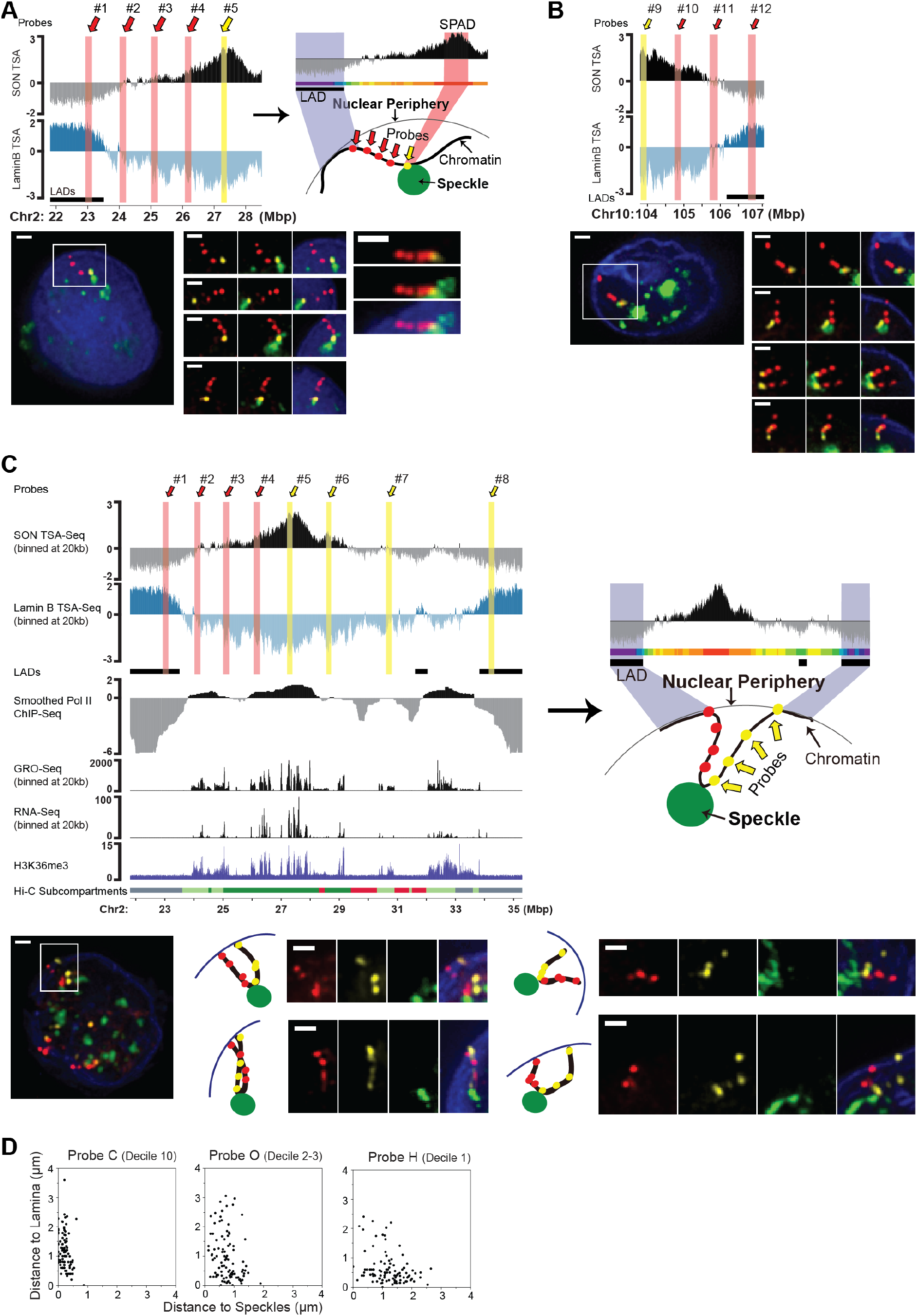
TSA-Seq predicts chromosome trajectories between nuclear periphery and speckle. (A) Immuno-FISH against 4 Mbp trajectory connecting SON TSA-Seq valley and peak. Top left: SON and Lamin B TSA-Seq genomic plot and FISH probe locations. Top right: schematic of TSA-Seq signal versus chromosome trajectory. Bottom: Probes 1-4 (red) and probe 5 (yellow) FISH and DAPI (blue). Left-Boxed region over FISH nuclear signal; Middle- (top) trajectory viewed in different channels and merged image; (lower) 3 additional examples. Right- 5th example. Scale bar: 1μm. (B) Immuno-FISH against 3 Mbp trajectory connecting SON TSA-Seq peak and valley. Top: TSA-Seq genomic plots and FISH probe locations. Bottom left: Probe 9 (yellow) and probes 10-12 (red) FISH and lamin B (blue). Boxed region over FISH nuclear signal. Bottom right: boxed region viewed in different channels (top panel), plus 3 additional examples (lower panels). Scale bars: 1μm. (C) Immuno-FISH against ~ 11 Mbp trajectory from SON TSA-Seq valley to peak to valley. Top left panel (top to bottom): probe locations and SON TSA-Seq, lamin B TSA-Seq, smoothed, normalized RNA pol II ChIP-Seq, RNA-Seq, GRO-Seq (20 kb bins), H3K36me3 ChIP, and Hi-C subcompartments. Right: schematic of TSA-Seq signal versus chromosome trajectory. Bottom: Probes 1-4 (red), probes 5-8 (yellow) FISH over lamin B (blue) staining. Left-Boxed region over FISH nuclear signal. Middle and right: different channels (top, middle) plus 3 additional example. Scale bars: 1μm. (D) 2D scatter-plots of distances (μm) from nuclear speckle (x-axis) and periphery (y-axis) for FISH probes C (decile 10), O (deciles 2-3), and I (decile 1) (n=100).

In each of these trajectories, the linear slopes seen in both the SON versus lamin TSASeq data appear highly inversely correlated, consistent with TSA-Seq reporting distance from the target compartments. These linear slopes contrast with various readouts of transcription that vary significantly along these same trajectories (Figure 4C and S4C), indicating any small fraction of SON protein associating with nascent transcripts must be too small to contribute significantly to the TSA-Seq signal, consistent with our previous demonstration of using the TSA signal exponential decay to accurately predict the mean speckle distance of chromosome regions (Figure 2F-G).

### Exceptionally deterministic positioning of specific chromosome regions at nuclear speckles

The radial positioning of different chromosome regions is thought to be stochastic (Bickmore, 2013; Takizawa et al., 2008). In contrast, using DNA FISH we observed that ~100% of the alleles from 3 different chromosome regions being probed, all belonging to the SON TSASeq top 5% percentile, map within 500 nm to nuclear speckles (Figures 2E, S2B). This level of deterministic positioning of a chromosome region relative to a defined nuclear compartment is exceptional. We thus define chromosome regions with SON TSA-Seq scores in the top 5% as “Speckle Associated Domains” (SPADs), which have a calculated mean distance to nuclear speckles of less than 0.32 μm (Figure 2F-G, S2). Extrapolating from our FISH results, we predict most other SPADs would also map deterministically near nuclear speckles.

### Nuclear lamina and speckles as “anchor points” for chromosome nuclear positioning

Single-cell FISH analysis reveals that SPADs are deterministically localized near speckles, but show large variability in nuclear lamina distance (Figures 4D, S5, probe C). Conversely, probes from SON TSA-Seq valleys (lamin TSA-Seq peaks) showed a high fraction of alleles positioned near the nuclear lamina but broad nuclear speckle distance distributions (Figure 4D and S5, probe R). Probes neither close to lamina nor close to speckles have broad distributions in both speckle and lamina distances, as observed in single cells by DNA FISH. The variable distances of SPADs to the lamina and LADs to speckles reflects the variable intranuclear spatial distribution of speckles. Thus the nuclear lamina and speckles act to “anchor” LADs and SPADs, respectively, with variable radial positioning of intermediate sequences. Instead, the linear nuclear lamina-speckle axis revealed by 2D scatter-plots of lamina versus speckle TSA-Seq represents an ensemble average over the cell population.

### Variation in genomic and epigenomic features as a function of speckle and lamin TSA-Seq scores

We next examined how genomic and epigenomic features distribute with respect to relative distances from nuclear speckles and lamin as estimated by TSA-Seq. We first calculated histograms showing segregation of different features as a function of SON TSA-Seq deciles, with the 1^st^ and 10^th^ deciles corresponding to the 10% lowest and highest scores, respectively (Figures 5A-5C, S6).

**Figure 5.**
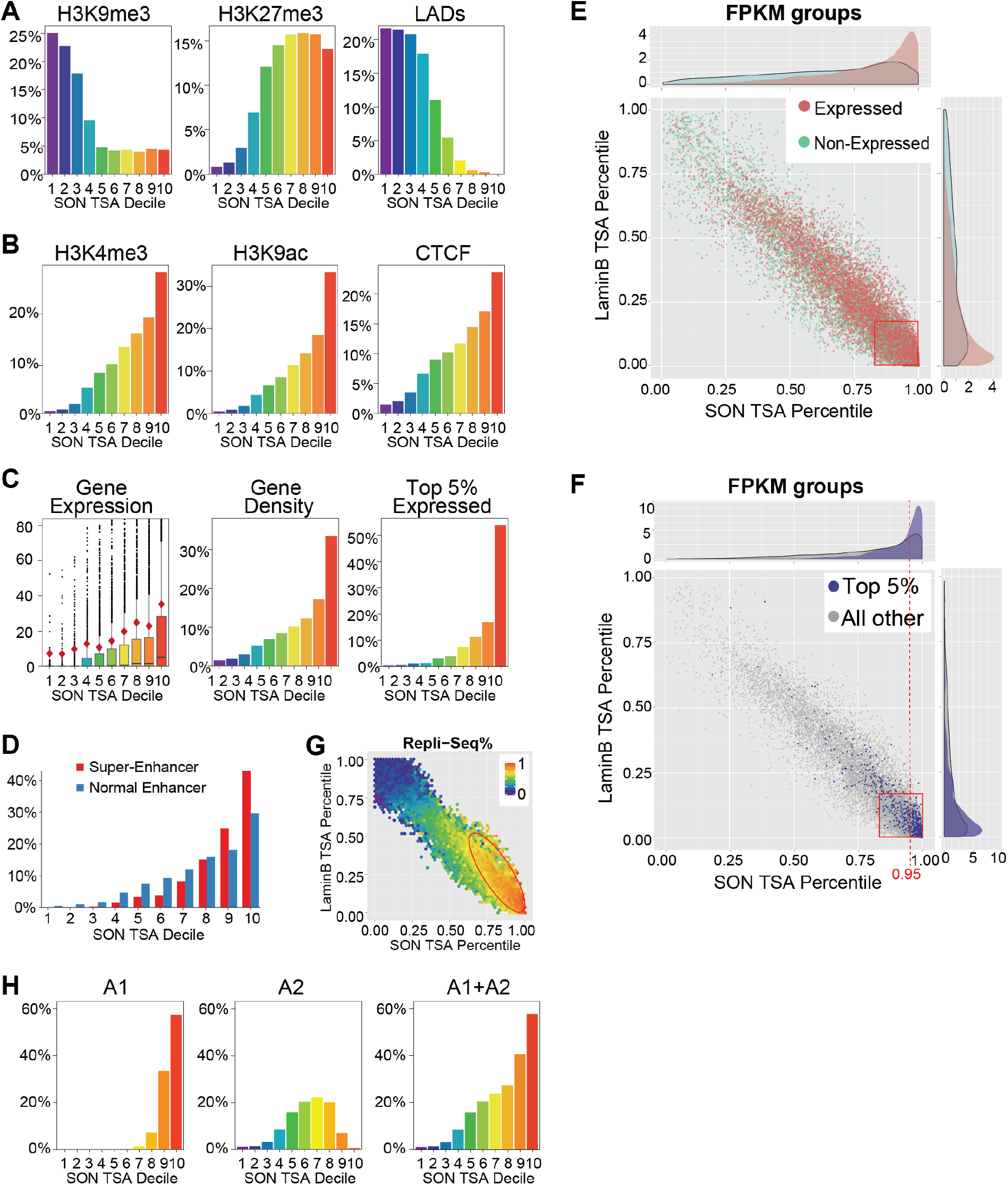
Gradients in transcription and DNA replication timing along speckle-lamin TSASeq axis reflect ensemble averaging over cells and chromosome regions. (A-D) gene expression and chromatin marks as function of SON TSA-seq deciles. (A) H3K9me3, H3K27me3, and LAD fractions. (B) H3K4me3 (peaks), H3K9ac, and CTCF (peaks). (C) Protein-coding gene expression (FPKM) (left), gene density (middle), and % of top 5% active genes (right). Box plot (left) shows median (black line), mean (red diamond), 75% (box-top), whiskers equal to 1.5x box size, and outliers (black dots). (D) Super-enhancers versus enhancers (%). (E-F) Lamin B versus SON TSA-Seq percentile 2D scatterplots (projections at top and right): (E) expressed (red) versus non-expressed (green) protein-coding genes. (F) top 5% expressed genes (blue) versus remaining expressed protein-coding genes (grey). Top 5% expressed genes furthest from nuclear lamina (right, red box) show disproportionate skewing towards speckles (> 95% SON TSA-Seq percentile, dashed line). (G) DNA Replication timing (160 kb bin size): earliest to latest (red to blue). Red ellipse-slight skewing of early replicating regions towards higher SON TSA versus lower lamin B TSA percentiles. (H) Distributions of A1 (left), A2 (middle), and A1 plus A2 (right) Hi-C subcompartments as a function of SON TSASeq deciles.

We found a roughly binary division for heterochromatin-related marks: LADs and H3K9me3 are enriched in low SON TSA-Seq deciles and depleted in high SON TSA-Seq deciles; H3K27me3, marking facultative heterochromatin, showed the inverse binary relationship (Figure 5A). H4K20me1, alternatively associated with silent (Sims et al., 2006) versus active chromatin (Wang et al., 2008), also shows a binary pattern similar to H3K27me3 (Figure S6C).)

In contrast, nearly all features associated with gene expression showed a linear or steeper gradient spanning all SON TSA-Seq deciles (Figures 5B, 5C and S6C). This includes enhancers, with super-enhancers showing a noticeably steeper gradient than normal enhancers (Figure 5D), indicating enrichment of super-enhancers at the speckle periphery. Gene expression and gene density, accompanied by decreased gene length, also show progressive increases with SON TSASeq decile (Figures 5C, S6A and S6B).

2D scatterplots show that genes with higher expression levels typically show both higher SON and lower lamin TSA-Seq percentiles (Figures 5E, 5F and S7A), while genes with intermediate expression levels show broader distributions along this lamin-SON TSA-Seq axis (Figures 5E and S7A). The most highly expressed genes show a pronounced shift away from the nuclear lamina and towards nuclear speckles (Figures 5C, 5F, and S7A), with >50% of the top 5% expressing genes mapping to the top SON TSA-Seq decile (Figure 5C, right); moreover, their distribution is disproportionally skewed towards being closer to nuclear speckles as compared to furthest from the nuclear lamina, particularly for the SPADs (Figures 5F, dashed line). Thus a closer mean distance to speckles, rather than a larger distance from the lamina, better describes the spatial positioning of the most highly expressed genes.

Similarly, DNA replication timing also distributes along this 2D linear speckle-lamin axis: The earliest replicating chromosome regions cluster nearly uniformly towards lower lamin and higher SON TSA-Seq scores (Figures 5G and S7B), with only a slight skewing of earlier replicating regions towards disproportionally higher SON TSA-Seq percentiles and intermediate replicating regions towards lower lamin B TSA-Seq percentiles (Figures 5G and S7B, red ellipses).

Notably, the observed linear gradient in transcription features as a function of distance to nuclear speckles (Figure 5B-D) represents an ensemble averaging over different and noncontiguous chromosome regions. For example, the observed gradient in transcription in the 1D SON TSA decile plot can be deconvolved as a superposition of the distributions of A1 and A2 active Hi-C subcompartment regions (Figure 5H).

### A subset of transcriptionally active “hot-zones” map close to nuclear speckles

Further comparison of SON TSA-Seq and transcription-related features in the context of chromosome neighborhoods reveals two types of “transcription hot-zones” (Figure 6A). We define “transcription hot-zones” as Mbp-sized regions with high RNA pol II occupancy and transcription, where pol II occupancy is measured by a smoothed and normalized RNA pol II ChIP-Seq signal (see Methods), and transcription is measured by GRO-Seq, RNA-Seq and H3K36me3 ChIP-Seq. These transcription hot-zones are also high in gene density. Strikingly, each SON TSA-Seq local maximum localizes near the center of each of the transcription hot-zones (red tick marks, Figure 6A). We define these SON TSA-Seq local maxima as the transcription hot-zone “peaks”.

**Figure 6.**
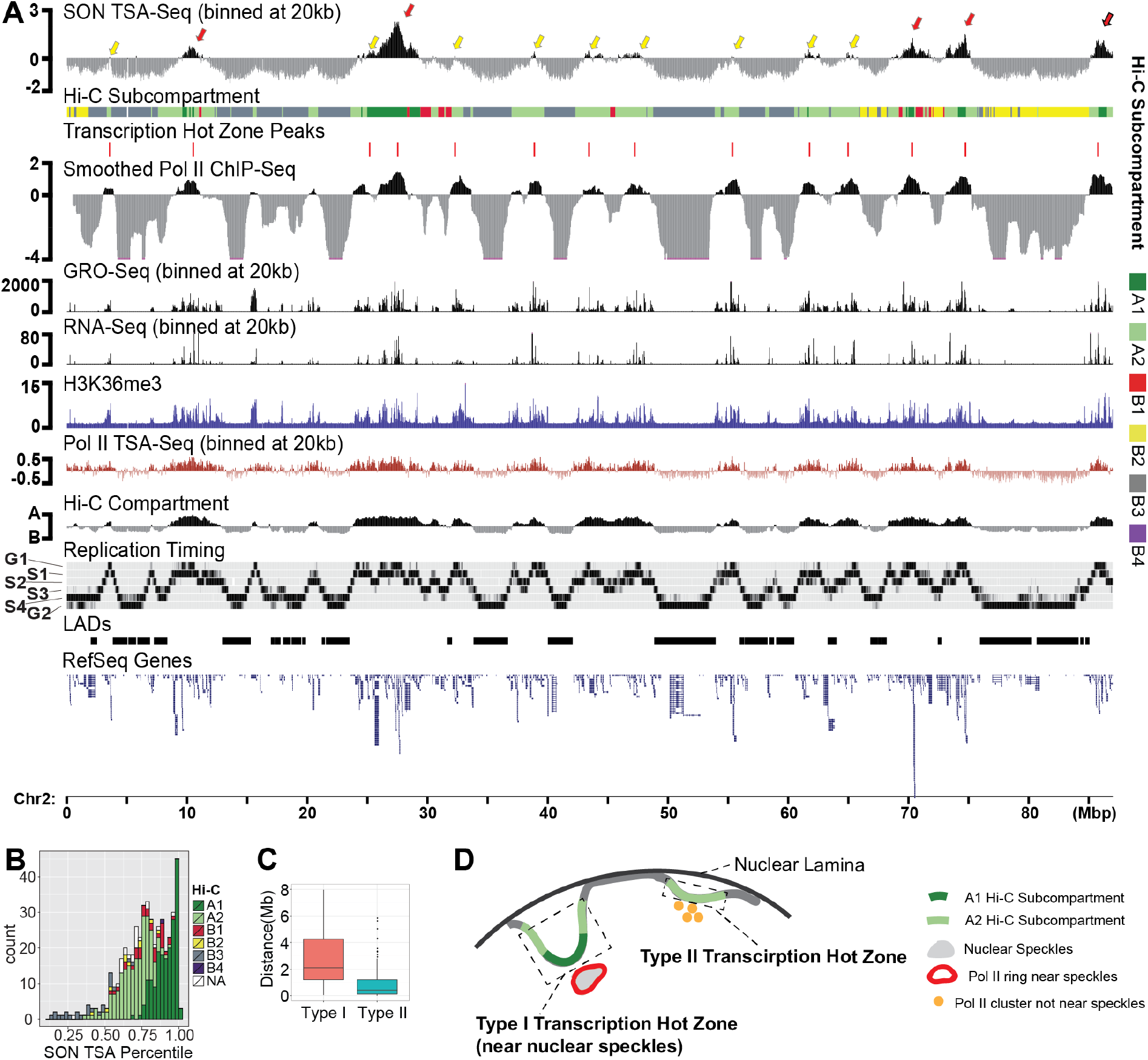
Two types of transcription “hot-zones” align with TSA-Seq local maxima. (A) Peaks of ~Mbp transcription “hot-zones”, marked by smoothed RNA pol II ChIP-Seq maxima align with SON TSA-Seq maxima (red tick marks). Top to bottom (Chr2): SON TSA-Seq, Hi-C subcompartments, smoothed, normalized RNA pol II ChIP-Seq, GRO-Seq (20 kb bins), RNASeq, H3K36me3, RNA pol II TSA-Seq, Hi-C eigenvector (compartment), Repli-Seq, LADs, RefSeq gene annotations. Arrows showing SON TSA-Seq local maxima align with type I (Red) versus Type II (yellow) transcription hot-zones. (B) Bimodal distribution of SON TSA-Seq local maxima number (y-axis), color-coded by A1 (dark green) versus A2 (light green) Hi-C subcompartment identity, versus SON TSA-Seq percentile (x-axis). (C) Box plots of genomic distances between SON TSA-Seq local maxima and nearest LAD / inter-LAD boundary for A1 (red) versus A2 (blue) SON TSA-Seq peaks. (D) Two types of transcription hot-zones: Type I whose apexes map adjacent to nuclear speckles and Type II whose apexes map to intermediate distances from nuclear speckles.

Transcription hot-zone peaks show a noticeably larger variation in the magnitude of the SON TSA-Seq peak value than the variation in magnitude seen for peaks of either the RNA pol II ChIP-Seq or RNA pol II TSA-Seq signals. The centers of transcription hot-zones containing the largest SON TSA-Seq peaks appeared to map to A1 Hi-C subcompartments (Figure 6A, red arrows), while the centers of transcription hot-zones containing smaller SON TSA-Seq local maxima appeared to map to A2 Hi-C subcompartments.

We sorted transcription hot-zone peaks according to their SON TSA-Seq percentile, creating a histogram of transcription hot-zone peak amplitudes (Figure 6B). The observed bimodal distribution suggests the existence of two classes of transcription hot-zones: Type I contains large SON TSA-Seq peaks, centered over A1 Hi-C subcompartment regions, which are typically flanked by a subclass of A2 Hi-C subcompartment regions (Figure 6A, red arrows). Type II contains smaller SON TSA-Seq peaks and peaks within valleys, centered over A2 Hi-C subcompartment regions, which are flanked on both sides by repressive Hi-C subcompartment regions (Figure 6A, yellow arrows).

Topologically, the SON TSA-Seq local maxima correspond to the most interiorly located regions of a local chromosome trajectory, as speckle and lamin TSA-Seq scores are inversely correlated (Figure 3). Mean genomic distances between the peaks of transcription hot-zones and their nearest LAD are larger for Type I versus type II transcription hot-zones (Figure 6C). These distance distributions suggest a degree of genomic “hard-wiring” of these two types of transcription hot-zones contained within simple inter-LADs (Figure 6D): Type I transcription hot-zones span A1 subcompartment regions and extend into flanking A2 subcompartment regions, are longer in length, protrude further into the nuclear interior, and have apexes predicted by their large SON TSA-Seq peak scores to localize adjacent to nuclear speckles. Type II transcription hot-zones span single A2 chromatin domains, are shorter in length, protrude smaller distances into the nuclear interior, and have apexes predicted by their smaller SON TSA-Seq peak scores to localize at intermediate distances to nuclear speckles.

### Transcription hot-zones near nuclear speckles have higher overall gene density, significantly higher number of the most highly expressed genes, and are enriched in housekeeping genes and genes with low transcriptional-pausing

Besides their different spatial localization relative to the nuclear lamina and speckles, what other differences might distinguish these two types of transcription hot-zones? We plotted the average gene density as a function of genomic distance from peaks of transcription hot-zones (Figure 7A). Total gene densities and the top 10% expressed gene densities are ~2-2.5-fold higher over Type I versus Type II transcription hot-zones. Both types of transcription hot-zones have ~2-fold higher densities of the top 10% expressed genes within 100 kb versus >200 kb of their peak locations. Whereas Type I transcription hot-zones also show total gene density and the density of moderately expressed genes peaking near their centers, Type II transcription hot-zones have near constant total gene and moderately expressed gene densities over the 500 kb regions flanking their peaks. These results indicate that transcription hot-zones near nuclear speckles have higher overall gene density, significantly higher number of the most highly expressed genes.

**Figure 7.**
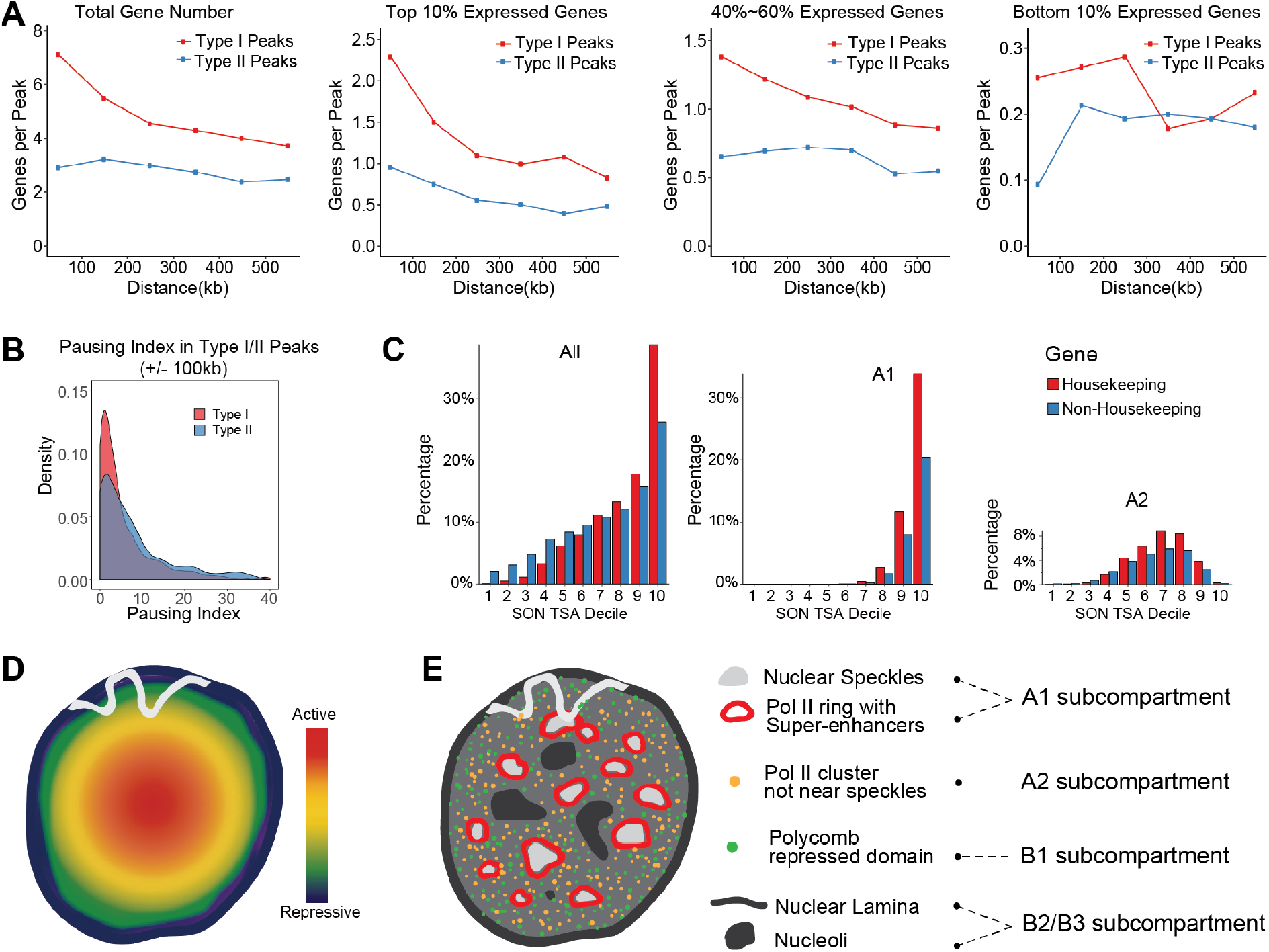
Differences between Type I and II transcription hot-zones and a refined model for nuclear organization. (A) Left to right: density (y-axis, gene number) of total genes, top 10% expressed genes, genes with 40-60% expression levels, and genes with 10% lowest expression as function of genomic distance (x-axis, kb) from centers of Type I (red) versus Type II (blue) transcription hot zones. (B) Pausing index distribution for expressed genes within +/- 100 kb of SON TSA-Seq local maxima contained within Type I (A1 peaks) versus Type II (A2 peaks) hot-zones. (C) Percentage (y-axis) of housekeeping versus non-housekeeping genes versus SON TSA-Seq deciles for all genes (left), genes only in A1 subcompartment regions near speckles (middle), or only in A2 subcompartment regions (right). (D) Previous model: transcriptional activity increases radially from low near the nuclear periphery to high towards the nuclear center. (E) New, refined model: Transcriptionally active chromosome regions are depleted from heterochromatic compartments close to the nuclear and nucleolar periphery (black). Apexes of chromosome trajectories protruding into nuclear interior correspond to centers of transcription hot-zones which localize either at the periphery (red rings) of nuclear speckles (grey) (Type I) or at interior sites at intermediate distances from speckles, possibly near unknown compartment such as transcription factories (Type II). A second repressive compartment, enriched in H3K27me3 (green dots), maps throughout the nuclear interior, dispersed between RNA pol II foci and speckles. Large-scale chromatin fibers (gray curves) traverse between these compartments.

We did not discover any GO terms special for genes near nuclear speckles (data not shown). However, we do see a significant enrichment of genes with low RNA pol II pausing index for speckle-associated Type I transcription hot-zones within 100 kb of their peaks (Figure 7B). We also see a relative enrichment of housekeeping versus non-housekeeping genes close to nuclear speckles (Figure 7C).

## Discussion

Here we presented TSA-Seq, the first genome-wide method capable of directly and quantitatively estimating mean cytological distances from particular nuclear subcompartments and even inferring local chromosome trajectories within the nucleus. TSA-Seq is fundamentally different from molecular proximity methods such as ChIP-Seq and DamID, that report only on the small fraction of the target protein in close proximity to DNA but are relatively insensitive to much larger concentrations of the same target protein that are not in direct molecular contact distance of the DNA. TSA-Seq instead converts an exponential decay of tyramide free radicals into a cytological “ruler” which can be used to probe relative distances over an ~1-1.5 μm radius from the staining target. In the case of nuclear speckle TSA staining, we demonstrated the capability of converting TSA-Seq scores into actual mean distances with an estimated accuracy of less than 100 nm.

Previous low-throughput FISH studies had led to a model in which a gradient of increasing transcriptional potential extends from the nuclear periphery to nuclear center (Bickmore, 2013) (Figure 7D). However, intranuclear spatial positioning of specific genes appeared to be highly stochastic. We propose that previous descriptions of stochastic positioning of chromosome regions as a function of gene expression relative to a nuclear periphery to center axis (Figure 7D) conceal a more deterministic positioning of different genomic regions relative to specific nuclear compartments (Figure 7E). One such compartment is the nuclear periphery (van Steensel and Belmont, 2017). Here we show that nuclear speckles act as a second “anchor” to organize interphase chromosomal organization.

Our TSA-Seq results instead suggest a refined model of nuclear organization (Figure 7E). Inter-LADs vary in how far they protrude into the nuclear interior, with centers of transcriptional “hot-zones” enriched in the most highly expressed genes, mapping to the most interior location of an inter-LAD chromosome trajectory. A fraction (~5%) of the genome associated with nuclear speckles with ~100% probability, with mean distance to speckle periphery less than 0.32 um. Transcription hot-zones mapping closest to nuclear speckles correspond to the A1 Hi-C subcompartment, have ~2-fold higher gene densities, and are enriched in super-enhancers, the most highly expressed genes, and genes with low transcriptional pausing. Interestingly, factors involved in RNA pol II pause release (pTEFb, cdk12) are enriched in nuclear speckles (Dow et al., 2010; Herrmann and Mancini, 2001; Ko et al., 2001).

Both transcriptional activity and timing of DNA replication align along a nuclear speckle-lamina axis, which we show represents an ensemble average over different cells as well as different types of noncontiguous chromosome trajectories. Our model describes several spatially distinct nuclear compartments: a) a thin rim adjacent to the nuclear periphery and relatively depleted of RNA pol II and transcription and enriched in LAD regions, likely also localizing at the periphery of nucleoli and centromeres (van Steensel and Belmont, 2017), not directly probed in this study; b) interior regions enriched in A2 Hi-C active subcompartments; c) intermingled regions enriched in polycomb-silenced regions corresponding to the B1 Hi-C active subcompartment; and d) the periphery of nuclear speckles where A1, Type I transcription hot-zones terminate (Figure 7E). Our TSA-Seq results in fact assign the differential localization of the Hi-C active A1 and A2 subcompartments to speckle-associated versus interior nuclear locations.

The logic of this nuclear organization remains to be determined. Our model would suggest, however, that chromosome movements of just several hundred nm during cell differentiation or during a cell physiological response might have significant functional significance. For example, a movement of just several hundred nm from the nuclear periphery towards the nuclear interior to sites enriched in RNA pol II foci might be needed to significantly boost the transcription of a gene contained within a repressive chromatin domain. Similarly, a movement of just a few hundred nm from the nuclear interior to adjacent to a nuclear speckle may similarly produce a significant transcriptional enhancement. Further analysis to more fully exploit TSA-Seq distance mapping should provide further insights. Additional insights will come from extension of TSA-Seq to additional nuclear compartments and examination of changes in nuclear organization accompanying developmental and cell physiological transitions.

## METHODS

### TSA-Seq

*TSA Cell Labeling*: K562 cells were obtained from the ATCC and cultured according to ENCODE Consortium recommendations (http://genome.ucsc.edu/ENCODE/protocols/cell/human/K562_protocol.pdf). SON TSA labeling was done in independent cell batches. For each batch, cells were divided equally and stained according to four labeling protocols: “Condition 1”, “Condition 2”, “Condition 3”, and “No 1° control”. Identical staining conditions were used except that 50% sucrose was included in the TSA staining buffer for the “Condition 2” protocol, 0.75mM DTT in addition to 50% sucrose was added in the TSA staining buffer for the “Condition 3” protocol, and the anti-SON primary antibody was omitted for the “No 1° control” protocol. Conditions, cell numbers and yield for all SON, pSC35, lamin A/C, and lamin B TSA-Seq experiments are summarized in Table S1.

Cells were fixed by adding 10 ml of 8% paraformaldehyde (Sigma-Aldrich) freshly prepared in CMF-PBS (calcium, magnesium-free phosphate buffered saline) to 40 ml of cells in culture media, incubating for 20 min at room temperature (RT), and quenching aldehydes by addition of 5.6 ml of 1.25 M glycine (Fisher Scientific) and incubation for 5 min at RT. All incubation steps used a rotating platform to mix the cell suspension. Cells were then pelleted by centrifugation and resuspended in CMF-PBS buffer with 0.5% Triton-X100 (Sigma-Aldrich) for 30 min at RT. All centrifugations were for 5 min at 100-200g. Cells were centrifuged at RT and resuspended in 1.5% H_2_O_2_ (Fisher Scientific) in CMF-PBS for 1h at RT to quench endogenous peroxidases, followed by 3 washes at RT in PBST (0.1%Triton-X100/CMF-PBS) by repeated centrifugation and cell pellet resuspension.

Cells were resuspended in IgG Blocking Buffer (5% Normal Goat Serum (Sigma-Aldrich) in PBST) at a concentration of 10^7^ cells/ml for 1 hr, followed by centrifugation and resuspension in primary antibody solution (Rabbit-anti-SON primary antibodies (Ab1, Atlas antibodies cat # HPA023535) diluted 1:1000 or 1:2000 in IgG Blocking buffer, or rabbit polyclonal-anti-SON primary antibodies (Ab2) diluted 1:2000 in IgG Blocking buffer, or mouse-anti-phosphorylated SC35 primary antibodies (Sigma-Aldrich cat # S4045) diluted 1:500 in IgG Blocking buffer, or mouse monoclonal anti-lamin A/C primary antibody (Clone 5G4) diluted 1:1000 in IgG Blocking buffer, or mouse monoclonal anti-lamin B primary antibodies (Clone 2D8) diluted 1:1000 in IgG Blocking buffer) at 10^7^ cells/ml, or mouse monoclonal anti-RNA polymerase II primary antibodies (clone 4H8) (Millipore cat # 05-623) at 0.5 × 10^7^ cells/ml and incubated 20-24 hrs at 4°C. For the no primary control, cells were incubated in IgG Blocking buffer only. Cells were then washed 3x with PBST, resuspended in IgG Blocking buffer with secondary antibodies at 10^7^ cells/ml, and incubated 20-24 hrs at 4°C. Secondary antibodies (Jackson ImmunoResearch) used were HRP conjugated, goat-anti-rabbit secondary antibody diluted 1:1000 or HRP conjugated goat-anti-mouse secondary antibody diluted 1:200 or 1:1000.

Cells then were washed 3x in PBST and resuspended in reaction buffer A (CMF-PBS for Condition 1, or CMF-PBS with 50% sucrose for Condition 2, or CMF-PBS with 50% sucrose and 1.5mM DTT for Condition 3) at 2 × 10^7^ cells/ml. Equal volumes of buffer B (8mM tyramide-biotin stock solution diluted 1:5000 with 0.003% H_2_O_2_ in CMF-PBS for Condition 1, or CMF-PBS with 50% sucrose for Conditions 2 and 3) were then added. Cells were incubated for 10 min and then washed 3x in PBST at RT. Tyramide-biotin was prepared using tyramide hydrochloride (Sigma), EZ-link Sulfo-NHS-LC-Biotin (Pierce), dimethyl formamide (Sigma), and triethylamine (Sigma), as previously described (Hopman et al., 1998).

#### Monitoring TSA labeling by anti-biotin immunostaining

After each TSA labeling batch, one drop of the cell suspension was placed on a glass coverslip, air-dried, and fixed with 1.6% paraformaldehyde in CMF-PBS at RT for 15 min. After washing in PBST 3 x 5 min, coverslips were incubated 30 min at RT with Streptavidin-Alexa Fluor^®^ 594 (Invitrogen) diluted 1:200 in CMF-PBS, washed 3 x 5min at RT in PBST, and mounted in anti-fade media (0.3 μg/ml DAPI (Sigma-Aldrich) /10% wt/vol Mowiol 4–88 (EMD) / 1% wt/vol DABCO (Sigma-Aldrich) / 25% glycerol / 0.1 M Tris, pH 8.5).

#### DNA Isolation from Labeled Cells

Cell lysis and protease digestion were done at 55°C with constant mixing in 10mM Tris, 10mM EDTA, 0.5% SDS and 500μg/ml proteinase K (NEB) for 8-12 hrs (10^7^ cells/ml). NaCl was added for a final 0.2 M concentration and the lysate incubated 20-24 hrs at 65°C with constant mixing to reverse formaldehyde crosslinking. DNA was extracted 2x using an equal volume of phenol (Fisher)/chloroform/isoamyl alcohol (25:24:1) and 1x with an equal volume of chloroform/isoamyl alcohol (24:1). Tubes were kept at 55°C for 1h with the cap off to evaporate phenol residues, and then treated with 25 μg/ml RNaseA (Qiagen) at 37° C for 1hr and extracted again. DNA was then precipitated with 1/10 volume of 3M Na Acetate, pH 5.2, and 2 volumes of 100% EtOH at −20 °C for 20 min – 2 hrs.

#### DNA Fragmentation

DNA was dissolved in ddH_2_0 and sonicated to 200-800 bp with the majority of the fragments at ~500bp using a Branson 450 Digital Sonifier (with Model 102 Converter and microtip) or a Bioruptor (UCD-200 or Pico). For the Branson Sonifier, each sonication was done twice with 250 μl volume in 1.5ml Eppendorf tubes on ice for 20 sec (1 sec chase, 1 sec pulse, total 40s). For the Bioruptor, sonication was done with 30s on and 30s off but the number of cycles varied from tube to tube.

#### Monitoring Biotin Labeling of DNA by Dot Blot

DNA was spotted onto Nitrocellulose membrane (0.45 μm, Bio-Rad) and UV cross-linked to the membrane (0.24 joules, UV Stratalinker 1800, Stratagene). The membrane then was blocked in SuperBlock (TBS) Blocking Buffer (Pierce) with 0.05% Tween (Sigma-Aldrich) for 1 hr, incubated 12-16 hrs at 4 °C with Streptavidin-HRP (Invitrogen) diluted 1:2500 in the same blocking buffer, washed at RT with TBST (TBS/0.05% Tween), treated with SuperSignal West Pico chemiluminescent substrate (Thermo Scientific), and exposed with CL-XPosure Film (Thermo Scientific). For TSA labeling with a specific antibody, we estimate biotin labeling of ~ 0.0004-0.03 biotin per 10 kb, depending on the target and TSA condition; based on an average DNA fragment size of ~ 500 bp, this produced ~ 0.002-0.15 % of fragments labeled with biotin, assuming one biotin per fragment.

#### Affinity Purification of Biotinylated DNA

DNA derived from TSA staining batches that produced satisfactory TSA labeling, as verified by microscopy and dot blot analysis, was pooled for affinity purification. The required ratio of DNA and Dynabeads M-270 Streptavidin (Invitrogen) was estimated from the anti-biotin dot blot results and the known binding capacity of the beads. Beads were washed 3x in Wash and Binding (W&B) buffer (5mM Tris-Cl, pH7.5, 0.5mM EDTA, 1M NaCl) using a magnetic stand. 10% of input DNA was set aside. Remaining DNA was mixed with an equal volume of 2x W&B buffer. In the initial protocol, DNA samples in 4x beads’ original volume were mixed with the beads and nutated at RT for 0.5 hr. The beads were immobilized in a magnet, the solution removed, and an additional 4x original bead volume of DNA in W&B buffer was added. This was repeated until all DNA from each staining condition had been bound to the beads. The last incubation was extended for 8 hrs at 4 °C. Beads then were washed at RT 4x with 1x W&B buffer, 4x with TSE 500 (20 mM Tris pH 8.0, 1% Triton, 0.1% SDS, 2 mM EDTA, 500 mM NaCl), 4x with 1x W&B buffer, and 4x with TE buffer (10 mM Tris, 10 mM EDTA, pH 8.0). In a later modified protocol, DNA samples were mixed with the desired amount of beads all at once regardless of the volume of DNA solution, and the mixture was incubated with rotation at RT for 1h and then 4 °C for 8 hrs. Beads then were washed 5 times. For each wash, beads were first resuspended in 1ml 1x W&B buffer with Triton added at 0.05% final concentration, then incubated at 55 °C on heat block for 2 min, then nutated at 55 °C for 2 min. Beads were transferred to a new tube after each wash.

After the final wash, beads were resuspended in 10mM Tris, 10mM EDTA, 0.5% SDS, and 1mg/ml proteinase K at 600μl per 100μl of the original bead volume. Free biotin was added using 30μl of a 2 mM stock solution per 100μl of the original bead volume and beads digested overnight at 55°C with rotation. DNA was then extracted using phenol/chloroform as described previously and precipitated using 0.05 μg/ μl glycogen (Fermentas) as a carrier with 1/10 volume of Na Acetate and 2 volumes of 100% EtOH for 12-16 hr at −20 °C.

#### Library preparation and high-throughput DNA sequencing

For each of the TSA-Seq experiment staining conditions, input and Streptavidin pull-down DNA were used to make sequencing libraries, using the Kapa Hyper Prep Kit (Kapa Biosystems). DNA was blunt-ended, 3’-end A-tailed, and ligated to indexed adaptors with 6nt barcodes. PCR amplification selectively enriched for those fragments with adapters on both ends. Amplification was carried out for 8-16 cycles with the Kapa HiFi polymerase (Kapa Biosystems, Woburn, MA). An Agilent bioanalyzer DNA high-sensitivity chip (Agilent, Santa Clara, CA) was used to determine library fragment sizes. Library fragment concentrations were measured by qPCR on a BioRad CFX Connect Real-Time System (Bio-Rad Laboratories, Inc. CA) prior to pooling and sequencing. Libraries were pooled at equimolar concentration and sequenced for 101 cycles on an Illumina HiSeq2000, HiSeq2500 or HiSeq4000 using a HiSeq SBS sequencing kit, version 4. The raw .bcl files were converted into demultiplexed compressed fastq files using the bcl2fastq v1.8.2 and later v2.17.1.14 conversion software (Illumina). 27-81 million 100 nt reads were obtained for each sample, with average quality scores > 30. Library preparation and sequencing were done by the High-Throughput Sequencing and Genotyping Unit of the Roy J. Carver Biotechnology Center at UIUC.

### Lamin B1 DamID

DamID of lamin B1 was performed in K562 cells as described (Vogel et al., 2007; Meuleman et al., 2013), except that prior to the DpnI digestion 1.5μg genomic DNA was treated for 1 hr with 5 units of Antarctic phosphatase (NEB, cat # M0289S) in 20μl of 1X Antarctic Phosphatase Reaction Buffer (NEB) followed by ethanol precipitation. This additional step prevents ligation of the adaptor to DNA fragments originating from apoptotic cells. Samples from 2 independent biological replicates were hybridized to human genome tiling microarrays (Nimblegen HD2) which contained 2.1 million probes designed as previously reported (Meuleman et al., 2013).

### 3D Immuno-FISH

FISH probes were made from BACs (Invitrogen, See Table S2, SI for BAC list). Probes were labeled with biotin or DIG using nick translation (BioNick kit; Invitrogen) according to manufacturer’s instructions, with a minor change for digoxigenin (Dig) labeling, in which the kit’s 10X dNTP mix was replaced by 0.125 mM Dig-11-dUTP (Roche), 0.375 mM dTTP (NEB), 0.5 mM dATP (NEB), 0.5 mM dCTP (NEB), 0.5 mM dGTP (NEB), 500 mM Tris-HCl, pH8.0, 50 mM MgCl_2_ and 500 ug/ml BSA (NEB). Alternatively, DNA was end labeled using terminal transferase as described elsewhere (Dernburg, 2011) with the following reagents: terminal transferase and 5x buffer (Fermentas) and dATP (NEB) and biotin-14-dATP (Life Technologies) for biotin labeling or dTTP (NEB) and Dig-11-dUTP (Roche) for Dig labeling. Concentrations of labeled and unlabeled nucleotides were 54 μM and 108 μM, respectively, for a 25 μl reaction with 2 μg fragmented DNA.

K562 cells were plated on poly-L-lysine (MW 70,000-150,000, Sigma-Aldrich) coated Circes #1.5 glass coverslips (Fisher Scientific) in a 24-well plate at 0.3ml per well and 10^5^−6×10^5^ cells/ml in RPMI culture media plus 10% FBS (Sigma). Cells attached after 15 min at 37°C in a humidified incubator with 5% CO2. Cells were then permeabilized in 0.1%Triton in CMF-PBS for 1 min at RT, fixed with freshly prepared 3.6% paraformaldehyde (Sigma-Aldrich) in CMFPBS for 10 min at RT, and quenched with 0.5M glycine (Fisher Scientific) in CMF-PBS for 5 min at RT.

Our 3D immuno-FISH was modified from that previously described (Solovei and Cremer, 2010). For anti-SON and anti-lamin B immunostaining, cells were blocked in IgG Blocking buffer for 1hr at RT, incubated with rabbit-anti-SON antibodies diluted 1:1000 with or without mouse-anti-lamin B antibodies diluted 1:250 in IgG blocking buffer overnight at 4°C, washed with PBST 3 x 5 min at RT, incubated 1 hr at RT with FITC-labeled, secondary goat-anti-rabbit IgG (Jackson ImmunoResearch) diluted 1:500 with or without AMCA labeled, secondary goat-anti-mouse IgG (Jackson ImmunoResearch) diluted 1:50 in IgG blocking buffer, and washed 3×5 min with PBST at RT. Cells were then post-fixed with 3% paraformaldehyde in CMF-PBS for 10 min at RT, quenched with 0.5M glycine in PBS for 5 min at RT, and washed with PBST for 5min at RT.

Coverslips were incubated in 20% glycerol in CMF-PBS for 30-60 min at RT, frozen and thawed 6x using liquid nitrogen, washed 3 x 5 min with PBST, rinsed and incubated on ice with 0.1M HCl/0.7% Triton/2x SSC for 10 min, and then washed 3x 5 min with 2x SSC. FISH probes (100 ng for probes made by nick translation, 20 ng for probes made using terminal transferase) in 4 ul of hybridization buffer (10% dextran sulfate, 50% formamide, 2x SSC) were then added to coverslips and sealed onto glass slides with rubber cement. Samples were denatured on a hot plate at 76°C for 3 min and then hybridized 18-72 hrs in a humid chamber at 37°C.

Coverslips then were washed at 70°C in 0.4x SSC for 2 min and at RT in 4x SSC for 5 min. Alternatively, they were washed at 37°C for 3 x 5 min in 2x SSC, and then at 60 °C for 3 x 5 min in 0.1x SSC, and at RT in 4x SSC for 5 min. The biotin and DIG signals were detected by incubation for 40 mins to 2 hrs at RT in a humidified chamber with Streptavidin-Alexa Fluor^®^ 594 (Invitrogen) diluted 1:200 and/or mouse-anti-DIG-Alexa ^®^Fluor 647 (Jackson ImmunoResearch) diluted 1:100 in 4x SSC with 4% BSA (Sigma). Coverslips were washed in 4x SSC/0.1%Triton-X100 3x 5min and mounted in DAPI-containing, anti-fade mounting media.

### Microscopy

3D optical-section images were taken with 0.2 μm z-steps using an Applied Precision personal DeltaVision microscope system and Deltavision SoftWorx software (GE Healthcare) with a 60x, 1.4 NA objective lens and a Cool-SNAP HQ_2_ CCD camera. Images were deconvolved using an enhanced ratio, iterative constrained algorithm (Agard et al., 1989) within the Applied Precision SoftWorx software (GE Healthcare). For immunoFISH images, 3D distances were measured from the center of each FISH spot to the edge of its nearest nuclear speckle using the Standard Two-Point Measure tool in SoftWorx. 3D surface plots were generated using FIJI software (Schindelin et al., 2012) with the Spectrum LUT. All figure panel images were prepared using FIJI and Photoshop CS6 (Adobe, San Jose, CA). Image enlargement used bicubic interpolation in Photoshop CS6.

### Exponential Fitting of microscopy and TSA-Seq data

For microscopy images, the green and red channel pixel intensities along a line were exported using the line profiling function in FIJI software and plotted against distance using OriginPro 9.1 (OriginLab, Northampton, MA). We used OriginPro 9.1 to fit the TSA signal to the equation y=y0+A*exp(R0*x), where y is the TSA signal intensity, and x the distance to the intensity peak location, using the iterative Levenberg Marquardt algorithm. All fitting results have adjusted R-square > 0.87. OriginPro 9.1 was also used to fit the TSA-Seq genomic data using the same equation and settings as used for the microscopy images. Here y is the TSA enrichment ratio between pull-down and input, and x is the mean speckle distance measured from 100 alleles by 3D immunoFISH. 8 FISH probes (probes A to H) were used to do the fitting while 10 additional probes (probes I to R) were used to test the goodness of the fit. All fitting results have adjusted R-square > 0.97.

### Read Mapping and Processing

Raw sequencing reads were first mapped to the human genome (UCSC hg19) using Bowtie 2 version 2.0.2 (Langmead and Salzberg, 2012) with default parameters. Chromosome Y was excluded in the reference genome to reduce mapping bias since the K562 cell line was derived from a female. PCR duplicates reads were removed using rmdup command from Samtools version 0.1.19 (Li et al., 2009) with default parameters.

We then normalized and processed the TSA-Seq data using a sliding window approach as follows. First, the normalized TSA-Seq enrichment score was calculated as in Eq. 1, where *N*_*TSA*_ is the number of mapped reads in a 20 kb window in the pull-down sample and *T*_*TSA*_ is the total number of mapped reads of pull down sample (*N*_*input*_ and *T*_*input*_ are defined similarly for input data). This approach can be interpreted as the fold change between the pulldown sample and the input corrected by the total number of mapped reads in each sample. The window slides on the genome with a step size of 100 bp.

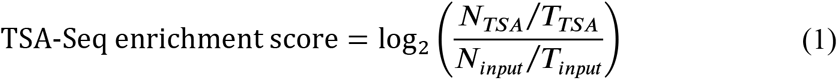

### SON TSA-Seq Correlations with other Genomic Features

We partitioned the genome into non-overlapping 20 kb bins, averaging the TSA-Seq score (Eq. 1) over the 100 bp intervals within this bin for the new TSA-Seq 20 kb score. The values for these 20 kb bins were then smoothed by convolution using a Hanning window of length 21 (21 x 20 kb or 420 kb). This new smoothed TSA-Seq score with 20 kb sampling was used for SON TSA-Seq decile calculations and comparisons with other genome features.

Various functional genomic data-sets available for K562 cells were downloaded from the ENCODE project portal (https://www.encodeproject.org/) (Table S3). For gene expression analysis, we used mRNA FPKM values based on RNA-Seq data for protein coding genes. Gene density was calculated by taking the fraction of protein coding genes in each SON TSA-Seq decile. For DNase I hypersensitive sites we calculated the number of DNase-Seq peaks contained within each decile. Histone modification and transcription factor ChIP-Seq data are classified into punctuated peaks, broad peaks, or a mixture of punctuated and broad peaks based on signal enrichment patterns. ChIP-Seq data (H3K27ac, H3K4me1, H3K4me2, H3K4me3, H3K79me2, H3K9ac, CTCF) with punctuated peak patterns show sharply defined peaks of localized enrichment. ChIP-Seq data (H2A.Z, H3K27me3, H3K36me3, H3K9me1, H3K9m3, RNA pol II) with broad peak patterns show diffuse, wide-spread genomic ChIP-Seq signals without discrete peaks. ChIP-Seq data with a mixture pattern of punctuated peaks and broad peak show both local and large-scale signal enrichment (H4K20me1).

For ChIP-Seq data with punctuated peaks, we counted peak number for each SON TSASeq decile. For ChIP-Seq data with broad peaks, we processed the ChIP-Seq data using a similar sliding window method as used for the TSA-Seq data (Eq. 2), using normalized ENCODE bigWig files for both the ChIP sample and input DNA. *N*_*ChIP*_ is the sum of signals in a 20 kb window for the ChIP sample. *N*_*input*_ is the sum of signals in a 20 kb window for the input DNA. *EN*_*input*_ is the expected value for a 20 kb window based on the total number of mapped reads and the total genome-mappable bp. *N*_*ChIP_norm*_ can be interpreted as the ChIP signal normalized by the input DNA signal. The mode of the normalized ChIP scores *N*_*ChIP_norm*_ was considered as the non-specific antibody-binding background level, which was subtracted from the normalized score in each 20 kb window to calculate the final ChIP-Seq enrichment score, *E*_*ChIP*_ (Eq. 3). The summed ChIP-Seq enrichment scores were calculated for each decile. An exception was H3K9me3 for which real signals were present throughout the genome. For this mark, the 0.5% percentile value was assumed as background and subtracted from the normalized score. For data with a mixture of sharp and broad peaks, we calculated the total number of base pairs within the ChIP-Seq peaks (peak length) in each SON TSA decile.

Genome GC-content data was downloaded from the UCSC Genome Browser and the GC fraction in each 20 kb bin calculated for each decile. We downloaded the RepeatMasker track for hg19 from the UCSC Genome Browser to calculate distributions of different repeat types in each decile. We calculated these distributions using coverage for LINE repeats but repeat count for the shorter Alu, SINE, simple, and satellite repeats.

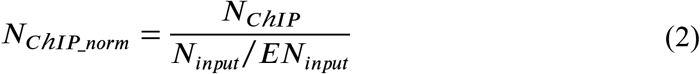

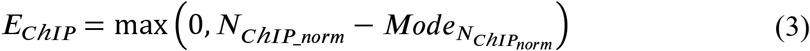

Replication timing and Hi-C subcompartment data from K562 and GM12878 cells, respectively, were compared with SON TSA-Seq deciles in two ways: 1. Calculating the fraction of each replication timing domain or Hi-C subcompartment group overlapping each SON TSASeq decile; 2. Calculating the fraction of each SON TSA-Seq decile that was comprised of each replication timing domain or each Hi-C subcompartment group. For display of replication timing, we also used the UW ENCODE Repli-Seq track corresponding to the wavelet-smoothed signal that combines data from different replication timing domains into one track.

For analysis of LAD distribution among the SON TSA-Seq deciles, we first followed the algorithm described previously for LAD segmentation (Guelen et al., 2008). Adjacent LADs were then merged if more than 80% of probes of the region between two LADs had positive log2 ratios.

For analysis of distribution of enhancers and super-enhancers across SON TSA-Seq deciles, we downloaded genomic locations of K562 super-enhancers from Hnisz, Denes, et al (Hnisz et al., 2013). The SON TSA decile assignment for each enhancer is picked by whichever has the largest fraction over the enhancer region.

### Box Plots

Each box plot shows median (inside line), 25th (box bottom) and 75th percentiles (box top). Upper whisker extends from 75th percentile to the highest value within 1.5-fold of the box length, and lower whisker extends from 25th percentile to the lowest value within 1.5-fold of the box length. Outliers are marked as black dots. Means are marked with red diamond, if shown.

### Quality Control with TSA-Seq

TSA-Seq is fundamentally different from molecular proximity methods such as ChIP-Seq. The amount of biotin-DNA labeling produced by the TSA reaction at a given chromosome locus is proportional to the intensity of the tyramide-biotin signal over that chromosome locus. Therefore the most essential control for TSA staining is direct microscopic visualization of the biotin intranuclear distribution after TSA staining. Labeling specificity should be validated by comparing the biotin staining with traditional immunofluorescence staining using the same primary antibody. TSA labeling should show both a specific labeling with a high intensity at the target compartment, as well as a diffuse signal surrounding the target which decays with distance from the target. We aliquoted cells for biotin staining and microscopic inspection after each batch of TSA staining. Batches showing unsatisfactory TSA staining were discarded. This was followed by dot blot analysis of biotin DNA labeling both before and after affinity purification. Again samples that did not show satisfactory biotin enrichment over negative control (no primary antibody) samples were discarded.

TSA-Seq is different because the tyramide free radical spreads over 100s of nm to a micron distance away from the staining target. ChIP-Seq results may vary between antibodies probing different components of a multi-subunit protein complex because of their varying distances to DNA and ability to cross-link to DNA. The target proteins themselves may show varying ability to cross-link to chromatin and survive the chromatin solubilization step of ChIP-Seq. In contrast, antibodies targeting different subunits of protein complexes should produce similar TSA-Seq results as long as the distance between these target subunits is small relative to the 10s-100s nm distance over which the concentration of tyramide free radical noticeably decays. Even antibodies targeting different protein complexes should produce similar TSA-Seq results as long as these different complexes are colocalized over a distance that is small relative to the microscopically observed distance over which the tyramide free-radical concentration noticeably decays. For example, for a decay constant of −4 (Condition 2), the TSA signal would drop only 4% at a distance of 10 nm from the target, 18% at a distance of 50 nm, and 33% at a distance of 100 nm.

Conversely, even ChIP-Seq validated antibodies may perform poorly with TSA-Seq. An antibody may show strong nonspecific or off-target binding without affecting ChIP results if the antibody is too far to effectively cross-link to DNA. Yet this nonspecifically or off-target bound antibody will produce a tyramide free radical signal which will diffuse and react with neighboring DNA and the amount of DNA reaction will be linearly proportional to the intensity of this nonspecific biotin signal as visualized by light microscopy.

### Comparison of Immunostaining with TSA labeling

CHO K1 cells were transfected with an EGFP-SON BAC transgene as described elsewhere^6^. Cells were cultured at 37°C with 5% CO2 in F12 media supplemented with 10% FBS (Sigma-Aldrich) and Zeocin (0.2ng/ml). Cells were fixed in 1.6% freshly prepared paraformaldehyde solution in CMF-PBS (calcium, magnesium-free phosphate buffered saline) for 15 min, and then permeabilized in 0.5% Triton-x100 (Sigma-Aldrich) in CMF-PBS for 30 min at RT. For immunostaining, cells were blocked in IgG Blocking buffer (CMF-PBS containing 0.1% Triton and 5% Normal Goat Serum (Sigma-Aldrich)) for an hour, incubated for 20-24 hrs at 4°C with primary antibody (Rabbit anti-SON (Atlas antibodies cat #HPA023535)) 1:2500 diluted in IgG Blocking buffer, and then secondary antibody (Goat anti-Rabbit-Texas Red, Jackson ImmunoResearch) 1:1000 diluted in IgG Blocking buffer for 20-24 hrs at 4°C.

TSA labeling was done as described above (TSA-Seq).

### Generation of polyclonal antibody against SON

Rabbit polyclonal antibody against hSON peptide (947-964, Cys-DPYRLGHDPYRLTPDPYR) was produced by Pacific Immunology Corp. Antigen peptide was synthesized, conjugated to carrier protein Keyhole Limpet Hemocyanin, and affinity purified for immunization. Antibody was affinity purified against the antigen peptide from the third bleed antiserum and stored in 50% glycerol solution at a concentration of 1.6 mg/mL.

### Generation of monoclonal antibodies against lamin A/C and lamin B

All of the ROD2 and tail domains of lamin A/C and lamin B (LB1 and LB2) were expressed in bacteria as GST fusion proteins. They were purified by FPLC and then mixed together prior to injection into mice for immunization. Western blotting was used to screen the monoclonal antibodies from hybridoma colonies for their lamin specificity. Monoclonal antibody from clone 5G4 showed lamin A/C staining with no cross-reactivity apparent at a dilution of 1/10000 and only trace reactivity at a dilution of 1/500. Monoclonal antibody from clone 2D8 showed lamin B staining with no cross-reactivity apparent at a dilution of 1/2000 and only trace reactivity to lamin A/C at lower dilutions. Dilutions of monoclonal antibodies for TSA staining described in Methods were of concentrated Bioreactor supernatants.

### Structured Illumination Microscopy (SIM)

3D SIM images were taken with 0.125 μm z-steps using an OMX Blaze V4 (GE Healthcare) with a 100x/1.4 NA oil immersion objective lens (Olympus), laser illumination (405nm, 488nm and 568nm), and Evolve 512 EMCCD camera (Photometrics). Solid state illumination was used for Cy5 in wide-field mode for the fourth channel (SON) in Figure 3C. Image reconstruction, channel registration and alignment were performed with Softworx software.

### Bioinformatics Analysis of Other Genome Features

For correlating SON TSA-Seq levels relative to DNA repeats, we recorded positions for different repeat classes/families using the UCSC RepeatMasker track. The GC content in each SON TSA-Seq decile was calculated using the nucBed program included in bedtools version 2.17 (Quinlan and Hall, 2010). The number of exon, intron, isoform and TSS were extracted from the GENCODE (Harrow et al., 2012) annotation file version 19 from the GENCODE website (http://www.gencodegenes.org/). Gene sizes and gene density across SON TSA-Seq deciles were extracted from the GENCODE annotation file version 3c, the same annotation file used for generating gene expression data.

### Hi-C Compartment Eigenvector

To generate the Hi-C compartment eigenvector track, we used a recently published high resolution Hi-C dataset^1^ from human K562 cells. First we downloaded the intrachromosome Hi-C raw contact matrices at 10 kb resolution and normalized the contact matrices using provided KR normalization vectors. Then we applied Principal Component Analysis (PCA), using the function “princomp” with default parameters in R, on the normalized contact matrix of each chromosome individually. As shown previously (Imakaev et al., 2012), eigenvector expansion is equivalent to PCA of the normalized contact matrix. The first principle component (PC1) corresponds to the eigenvector with the largest eigenvalue. Eigenvectors were normalized by magnitude. Here we only showed the first eigenvector.

### Applying the normalization method used for TSA-Seq to Pol II ChIP-Seq data

The normalization scheme used for TSA-Seq data-which computes relative enrichment/depletion over a certain bin size-can also be applied to ChIP-Seq data. We processed the ENCODE K562 Pol II ChIP-Seq and input raw fastq reads using the same alignment pipeline as used for the SON TSA-Seq data. Alignments from replicate ChIP-Seq experiments were merged. We calculated the normalized TSA-Seq enrichment score using Eq. 1. Here *N*_*Pol*2_ is the number of mapped reads in a 20 kb window in the RNA Pol II ChIP-Seq sample (*N*_*input*_ is defined similarly for input data). *B*_*Pol*2_ is the expected number of reads from the background nonspecific binding of RNA Pol II antibody to chromatin. To estimate *B*_*Pol*2_ we first ranked the genomic intergenic regions from longest to shortest. We used the top 5% longest intergenic regions, which we assumed to be gene deserts, to estimate background reads. For these largest intergenic regions, we calculated the average read count per base pair in the ChIP-Seq data. *B*_*Pol*2_ is the product of the size in bp of the sliding window with this average background read count per bp. The background adjustment for each bin is done by using the max(0, *N*_*Pol*2_ − *B*_*Pol*2_). Finally, *T*_*Pol*2_ is the total number of mapped read for ChIP-Seq sample after the background subtraction for each bin and *T*_*input*_ is the total number of mapped reads in input data.

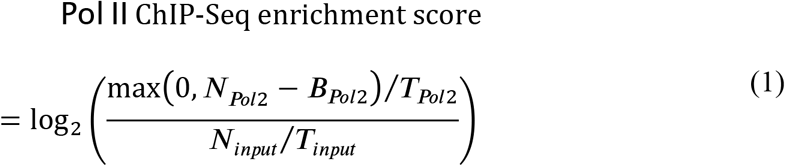

### Transcription hot-zones

To identify transcription hot-zones, we started with the smoothed RNA pol II ChIP-Seq track. We identified all genomic regions with positive values > 0.1, and for each of these regions we matched it to the local maximum in the SON TSA-Seq signal within this same region. If more than one local maximum was present within the region, we used the largest local maximum. In nearly all cases, these SON TSA-Seq local maxima aligned with the local maxima in the smoothed RNA pol II track. We identified the Hi-C subcompartment assignment corresponding to these SON TSA-Seq local maxima, as well as their SON TSA-Seq scores, and corresponding percentile. We then calculated the distance distribution for the A1 and A2 SON TSA-Seq local maxima to the nearest LAD / inter-LAD boundary.

### 2D TSA-Seq Scatter-plots

For 2D SON or Lamin B TSA-Seq and 1 Mbp-smoothed normalized RNA pol II ChIP-Seq scatter-plots (Figure S4C), each point represents a 160 kb bin for which the TSA-Seq average score was calculated from the smoothed TSA-Seq score. These points were plotted versus score percentile.

We also used similar 2D scatter-plots to show the relationship of SON, pSC35, lamin A/C, lamin B and Pol II TSA-Seq data in Figure 3B. Each point represents a 300 kb bin from the smoothed TSA-Seq score. These points were plotted versus score value or percentile. We also used these 2D scatter-plots to show the relationship between lamin and SON TSA-Seq with gene expression or DNA replication timing. In these scatter-plots (Figure 5E, 5F, 5G and S7), the x-y axes correspond to either TSA-Seq score values or percentiles of the TSA-Seq score over the entire genome, with distribution projections shown on the sides.

For gene expression, we calculated the FPKM (Fragments Per Kilobase of transcript per Million mapped reads) of each gene and divided genes into two groups-expressed (FPKM>0) and non-expressed (FPKM=0). For expressed genes, we further separated them into 10 deciles, ranking them based on their FPKM value. Each dot represents a single gene. For DNA replication timing, we used the UW Repli-Seq data for K562 cells (Hansen et al., 2010). Each dot represents a 300 kb genome bin. The fraction of each 300 kb bin replicating in each of the measured replication timing domains (G1, S1, S2, S3, and S4) was calculated and the bin was assigned to that replication domain with the largest fraction.

For computing the 2D TSA-Seq scatter-plot showing the distribution of Hi-C subcompartments (Figure 3F), we computed the average lamin B or SON TSA-Seq scores in 320 kb non-overlapping bins calculated from the smoothed 20 kb bin TSA-Seq scores as described above for the other 2D scatter-plots. For each 320 kb bin, the fraction of this bin corresponding to each Hi-C subcompartment was calculated. The Hi-C subcompartment with the largest fraction was then assigned to this 320 kb bin. TSA-Seq scores were plotted as percentiles.

### 2D TSA-Seq Plots of Weighted, Wavelet-smoothed DNA Replication Timing

We again used the UW replication timing data for K562 cells but now used the weighted, wavelet-smoothed composite signal which measures relative timing of DNA replication such that earlier replicating regions have higher values and ranked as percentiles. Average replication timing scores were calculated for 160 kb genomic bins. In Figure 5G, each 160 kb genomic bin was mapped to a given lamin B and SON TSA-Seq percentile based on the average TSA-Seq score calculated for each 160 kb bin using the smoothed TSA-Seq 20 kb bin data. We then divided the lamin B and SON TSA-Seq percentile ranges into hexagonal pixels spaced by 2 percentile. We calculated the median replication timing of all 160 kb genomic bins mapping to a given hexagon. These median values of replication percentiles were color-coded and plotted.

### Pausing Index of genes in TSA-Seq Peaks

We calculated the pausing index of each gene based on Pol II ChIP-Seq. The Pol II ChIP-Seq was downloaded from ENCODE (ENCSR000FAJ). The method of calculating pausing index was described in (Day et al., 2016).

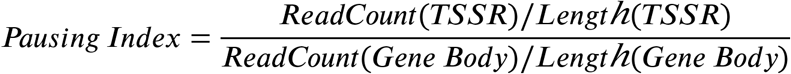

TSSR (TSS region, −50 to +300 bp around TSS). Gene body (TSS + 300 bp to +3 kb past the annotated transcriptional end site (TES).

Then we identified genes that were within the +/- 100kb range of TSA-Seq peaks. TSASeq peaks were separated into two groups based on whether they were in A1 or A2 subcompartments. In Figure 7D we plotted the density of the pausing index of genes near (+/- 200kb) A1/A2 TSA-Seq peaks.

### House-keeping genes in TSA-Seq Deciles

A list of 3904 human housekeeping genes was downloaded from http://www.tau.ac.il/~elieis/HKG/HK_genes.txt, which is an updated version of the original release from Eisenberg and Levanon, 2013. To determine the non-housekeeping genes, as well as the genomic position of these genes, we extracted protein-coding genes with completed CDS annotation from the hg19 RefSeq gene annotation (downloaded from the UCSC Genome Browser). 3791 genes among the original 3904 housekeeping genes were found to have matched gene id or gene name in the RefSeq gene annotation. Therefore, the remaining 15,751 protein-coding genes were determined as non-housekeeping genes. To compare the distribution of housekeeping genes to non-housekeeping genes across a combination of TSA-seq decile and Hi-C subcompartment, we determined the number of genes overlapping with a specific SON TSA-seq decile (e.g., decile 1) within a particular Hi-C subcompartment (e.g., A1). The number of overlapped genes were further normalized to a percentage for each type of Hi-C subcompartment.

## Data Availability

All high throughput sequencing data generated by TSA-Seq are deposited at GEO (http://www.ncbi.nlm.nih.gov/geo/) under GSE66019.

## Code Availability

All computational code used to analyze TSA-Seq data is available at: https://github.com/ma-compbio/TSA-Seq-toolkit

## Acknowledgements

This work was supported by National Institutes of Health grant R01 GM58460 to ASB, R01 HG007352 to JM, U54 DK107965 to ASB, JM, and BvS, and a NWO ZonMW-TOP grant to BvS. Sequencing was done at the UIUC Biotechnology Center. We thank William Brieher, Lisa Stubbs, and Brian Freeman (UIUC) for helpful suggestions during the course of this research.

## Author Contributions

YC developed the TSA staining and genomic mapping procedures, applied them to SON, pSC35, lamin A/C, lamin B, Pol II TSA-Seq genome mapping, processed sequencing data, and performed FISH validation experiments with guidance from ASB. LZ repeated SON TSA-Seq mapping with the original and a different SON antibody, and added SON Condition 3. LZ together with YC produced duplicates of the pSC35 and Pol II TSA-Seq and produced a duplicate of the lamin A/C TSA-Seq. Computational tool development and quantitative analysis of SON TSA-Seq data and integration with other genomic data was done by YZ and YW with guidance from JM and discussion among YC, ASB, YZ, YW, LZ, and JM. EKB and BvS generated lamin B1 Dam-ID genomic data from K562 cells. SA and RG provided monoclonal anti-lamin antibodies. Manuscript was written by YC and ASB.

---

**Table S1:** Summary of Conditions for all TSA-Seq experiments:

| Expt | Sample | Used for Figures | # of Cells | 1<sup>o</sup> Ab Conc. | 2<sup>o</sup> Ab Conc. | TSA Condi-tion | Round of Purifica-tion | Pull down amount | Round of PCR amplifi-cation |
| --- | --- | --- | --- | --- | --- | --- | --- | --- | --- |
| 1 | No primary control for SON TSA-Seq | Yes | 268 million | N/A | 1:1000 | TSA | 1 | 2.87 ng | 15 |
| 1 | SON TSA-Seq Ab1 Condition 1 | Yes | 268 million | 1:2000 | 1:1000 | TSA | 1 | 35.26 ng | 15 |
| 1 | SON TSA-Seq Ab1 Condition 1 2<sup>nd</sup> pull-down | Yes |  |  |  |  | started with 27.36 ng of input from the 1<sup>st</sup> pull down | 19.92 ng | 15 |
| 1 | SON TSA-Seq Ab1 Condition 2 | Yes | 268 million | 1:2000 | 1:1000 | TSA +Sucrose | 2 | 7.888 ng | 15 |
| 2 | SON TSA-Seq Ab1 Condition 1 Replicate | No | 415 million | 1:1000 | 1:1000 | TSA | 2 | 5.88 ng | 8 |
| 2 | SON TSA-Seq Ab1 Condition 2 Replicate | No | 415 million | 1:1000 | 1:1000 | TSA +Sucrose | 2 | 2.492 ng | 8 |
| 2 | SON TSA-Seq Ab1 Condition 3 | Yes | 590 million | 1:1000 | 1:1000 | TSA +Sucrose +DTT | 2 | ~1-1.5 ng | 8 for input; 12 for pull-down |
| 3 | SON TSA-Seq Ab2 Condition 2 | Yes | 270 million | 1:2000 | 1:1000 | TSA +Sucrose | 1 | 10.504 | 8 |
| 4 | No primary control for Lamin A/C TSA-Seq | No | 150 million | N/A | 1:200 | TSA | 2 | <1.122 ng | 8 for input; 10 for pull-down |
| 4 | Lamin A/C TSA-Seq Condition 2 | No | 150 million | 1:1000 | 1:200 | TSA +Sucrose | 2 | 3.78 ng | 8 |
| 5 | No primary control for Lamin A/C TSA-Seq Replicate | No | 710 million | 1:1000 | 1:1000 | TSA +Sucrose | 1 | 1.965 ng | 12 for input; 16 for pull-down |
| 5 | Lamin A/C TSA-Seq Condition 2 Replicate | Yes | 710 million | 1:1000 | 1:1000 | TSA +Sucrose | 1 | 23 ng | 12 |
| 6 | No primary control for Lamin B TSA-Seq | No | 400 million | N/A | 1:1000 | TSA +Sucrose | 1 | 1.364 ng | 10 |
| 6 | Lamin B TSA-Seq Condition 2 | Yes | 400 million | 1:1000 | 1:1000 | TSA +Sucrose | 1 | 4.8279 ng | 10 |
| 7 | No primary control for Lamin B TSA-Seq Replicate | No | 460 million | N/A | 1:1000 | TSA +Sucrose | 1 | 3.9 ng | 12 |
| 7 | Lamin B TSA-Seq Condition 2 Replicate | No | 460 million | 1:1000 | 1:1000 | TSA +Sucrose | 1 | 6.85 ng | 12 |
| 8 | pSC35 TSA Condition 1 | No | 485 million | 1:500 | 1:1000 | TSA | 1 | 32.136 | 8 |
| 9 | No primary control for pSC35 TSA-Seq Replicate | No | 326 million | N/A | 1:1000 | TSA | 1 | 3.435 ng | 12 |
| 9 | pSC35 TSA-Seq Condition 1 Replicate | Yes | 326 million | 1:500 | 1:1000 | TSA | 1 | 8.9 ng | 12 |
| 10 | Pol II TSA-Seq<br>Condition 1 | No | 225<br>million | 1:300 | 1:1000 | TSA | 1 | 52.44 ng | 8 |
| 11 | No primary<br>control for Pol<br>II TSA-Seq<br>Replicate | No | 175<br>million | N/A | 1:1000 | TSA | 1 | 2.3 ng | 10 for<br>input;<br>12 for<br>pull-down |
| 11 | Pol II TSA-Seq<br>Condition 1<br>Replicate | Yes | 175<br>million | 1:300 | 1:1000 | TSA | 1 | 14.05 ng | 10 |

**Table S2:**
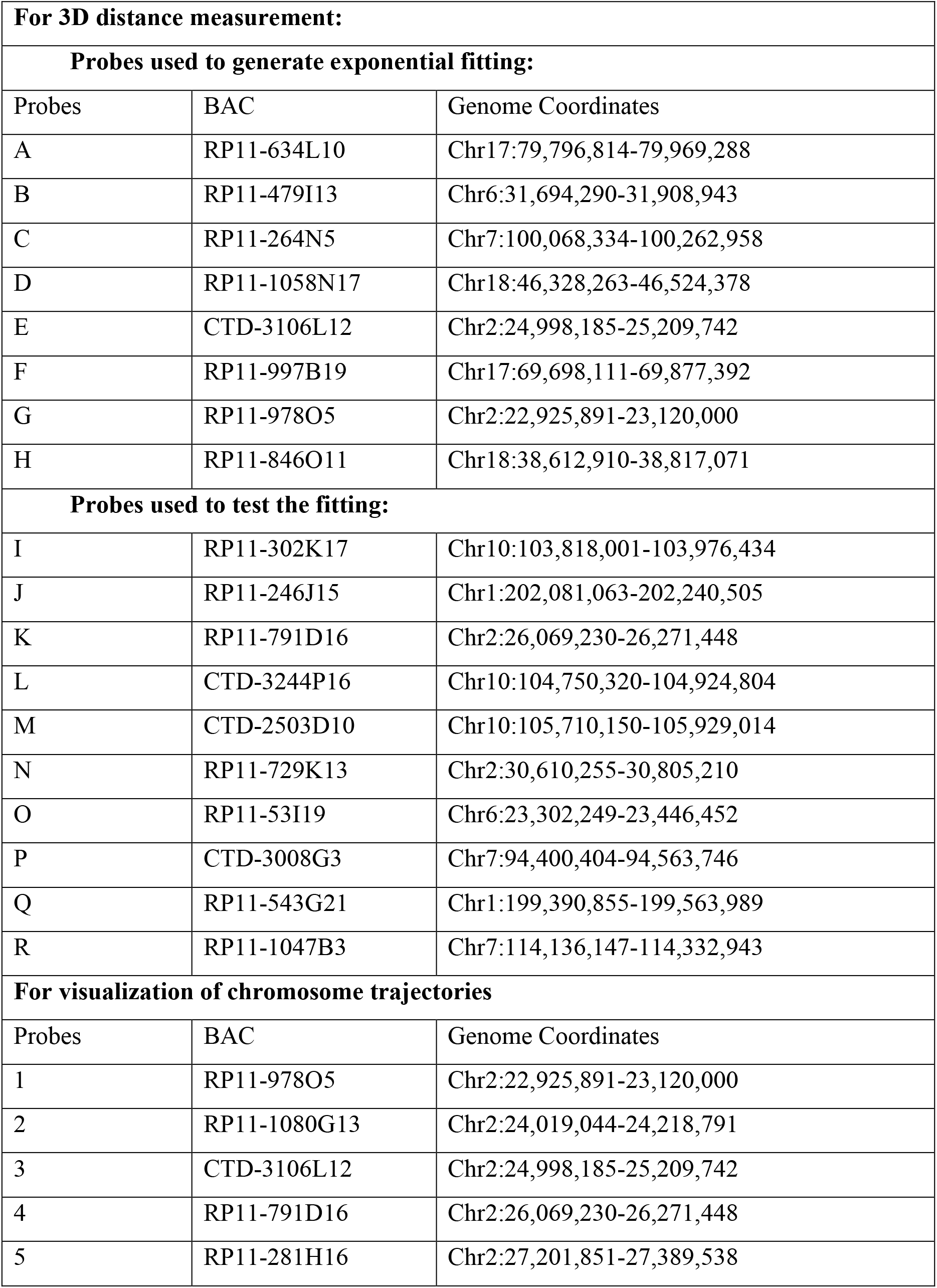
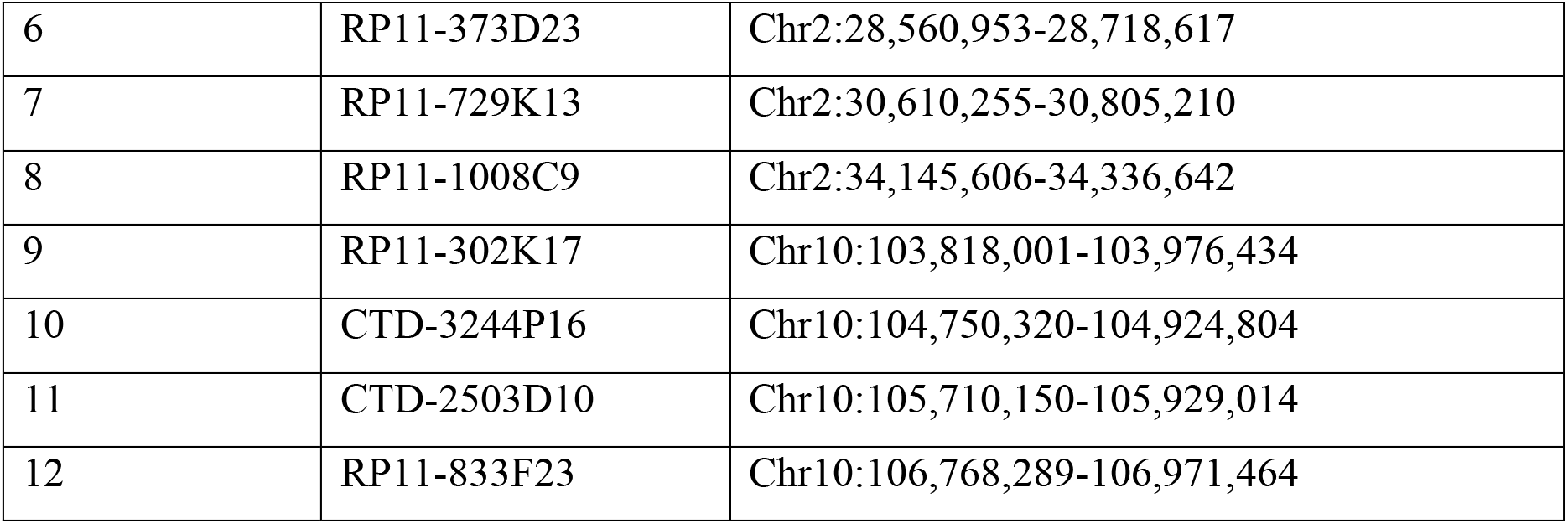
BACs used for DNA FISH

**Table S3: Public datasets analyzed.**

(See separate excel file)

**Figure S1.**
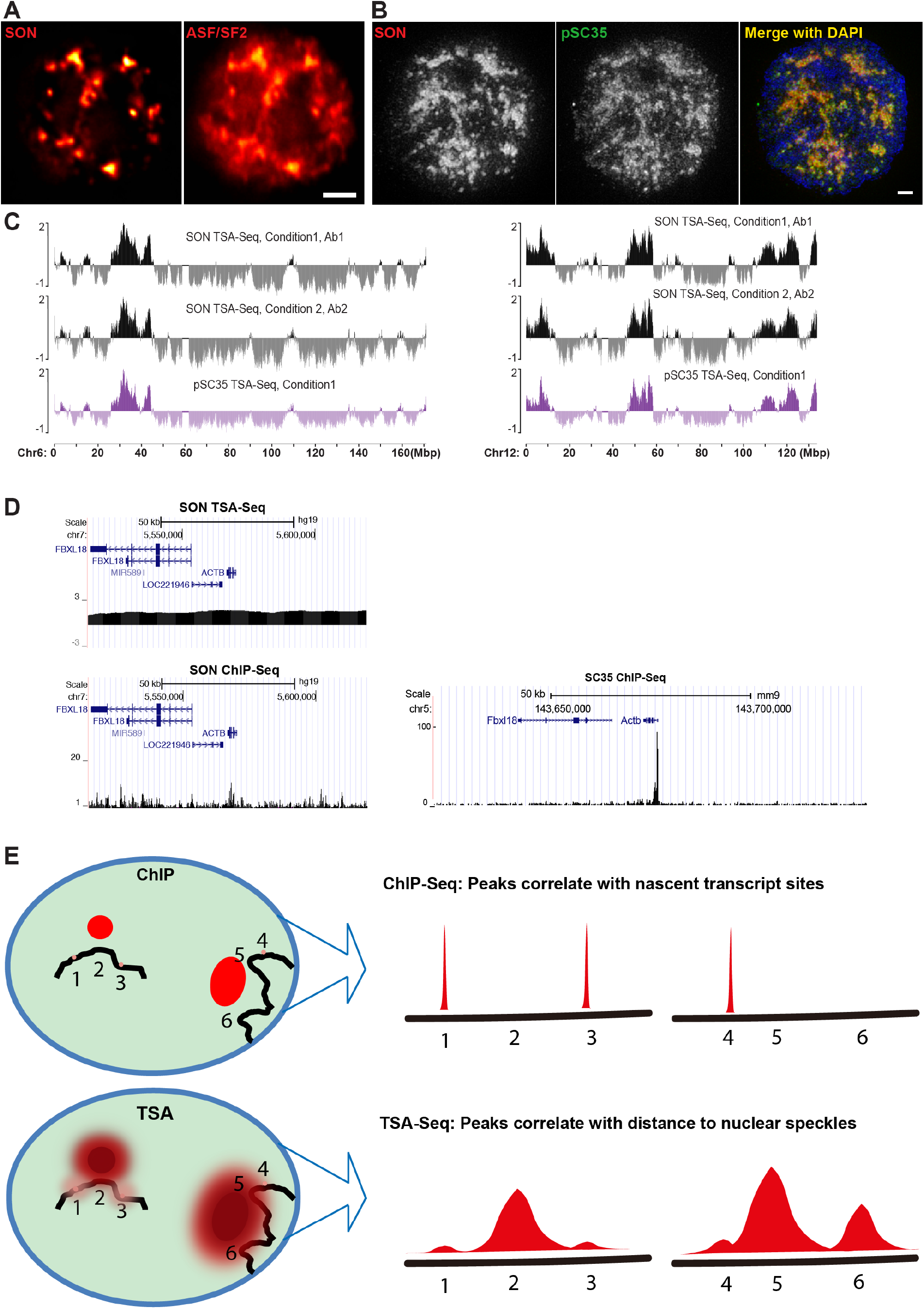
TSA-Seq measures cytological proximity. (A) SON immunostaining (left) provides much higher speckle to nucleoplasmic ratio than common speckle markers such as SR protein ASF/SF2 (right), as revealed by color heatmap display of wide-field light microscopy deconvolved optical sections. Scale bar: 2μm. (B) Left to right: 3D SIM imaging of SON, pSC35, and merged image with SON (red), pSC35 (green), and DAPI (blue). Images are projections from 15 z-sections with a 0.125 μm focus step size. Scale bar: 1μm. (C) Speckle TSA-Seq genomic plots generated with two different SON antibodies or pSC35 antibody for chromosome 6 (left) and chromosome 12 (right) all show near identical results: SON Ab1 (top) and Ab2 (middle), or pSC35 antibody (bottom). (D) Genomic plots comparing SON (Kim et al., 2016) or pSC35 (Ji et al., 2013) ChIP-Seq and TSA-Seq signals near ~100 kb region flanking ACTB gene in human (left, SON) or mouse (right, pSC35). (E) TSA-Seq is fundamentally different from ChIP-Seq: The TSA signal, generated by staining a protein concentrated in speckles (red ovals) spreads over micron distances (left), thereby generating a broad genomic signal (right), while ChIP requires molecular contact of protein with DNA, thereby producing a localized signal (right) only from protein present over specific DNA sequences (left, small red dots, sites 1,3, and 4). The speckle-derived TSA signal will dominate the overall TSA-Seq signal in proportion to the relative concentrations of the tyramide free radicals diffusing from the high concentration of speckle-localized protein versus from the small amount of localized SON protein near the DNA as visualized directly by light microscopy. Thus the SON TSA-Seq signal spans large genomic regions (D, left top), showing near constant levels over tens of kb regions.

**Figure S2.**
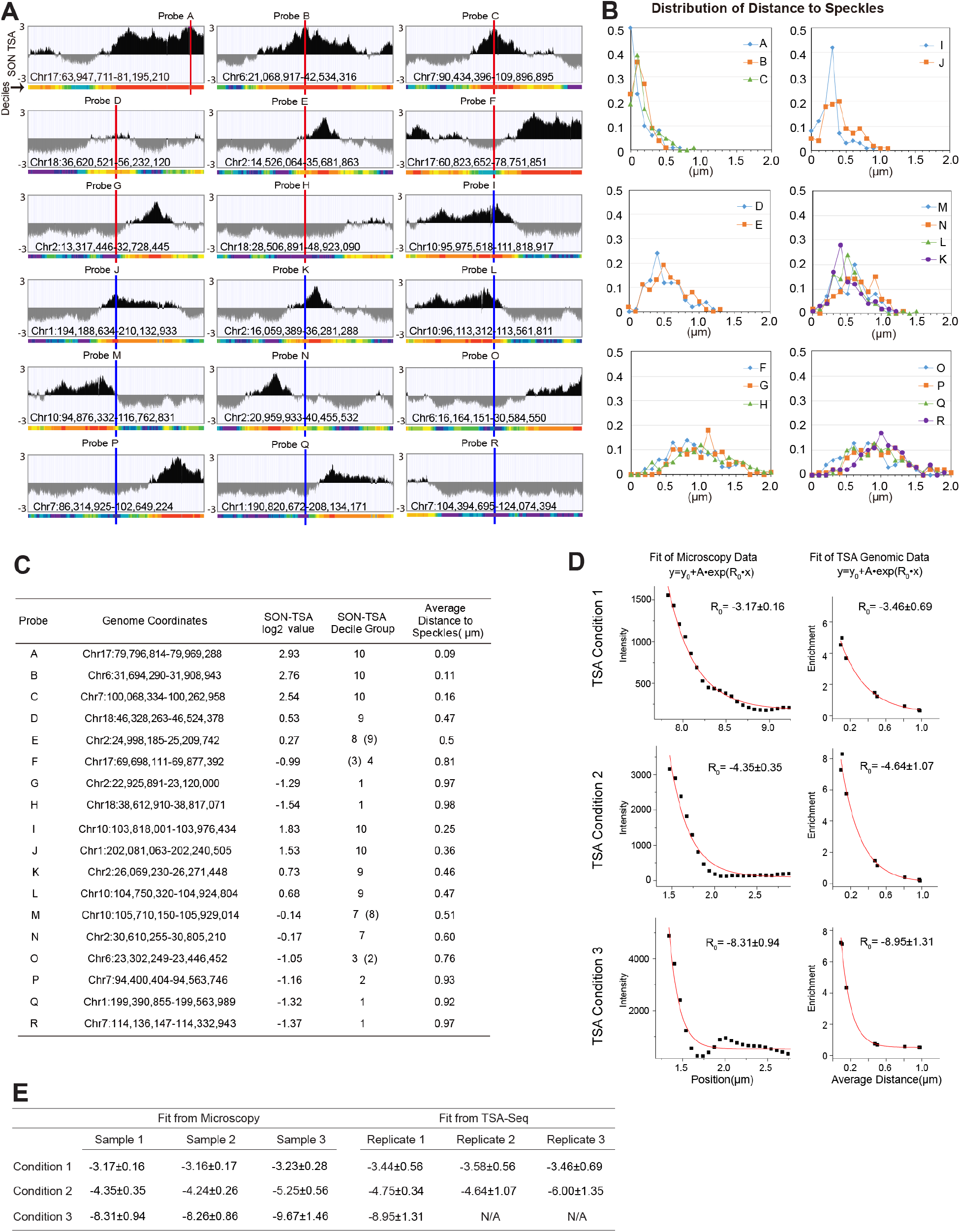
3D immuno-FISH validation that SON TSA-Seq signal measures cytological mean distance to nuclear speckles. (A) SON TSA-Seq genomic plots flanking 18 BAC probe locations (A-R), chosen from regions of different SON TSA-Seq scores and deciles. (B) Distance distribution to the nearest nuclear speckle for each of the 18 probes, as measured by 3D immuno-FISH (n=100). (C) Genome location of these 18 probes: SON TSA-Seq scores were averaged over the ~200 kb BAC FISH probe regions; TSA decile group shown corresponds to the majority of the probe region while value in parentheses shows decile for smaller part of BAC probe. Average distance is the mean of distance distributions shown in (B). (D) Exponential fitting of TSA microscopy (left) versus SON TSA-Seq (right) data for the three TSA staining conditions: Condition 1 (top), Condition 2 (middle), and Condition 3 (bottom). Left: lac repressor TSA-staining of integrated, multi-copy BAC array containing lac operator repeats (see text)-intensity versus distance from GFP spot. Right: exponential fit of SON TSA-Seq enrichment (relative to mean) versus mean speckle distance (C) for 8 genomic regions (probes A-H). (E) Fitting parameters and their standard error for TSA microscopy and SON TSA-Seq genomic data sets. The exponential decay constants from the direct cytological measurement of TSA spreading from lac repressor stained lac operator arrays are within experimental error of the decay constants measured from the fit to the TSA-Seq genomic data for each of the three TSA staining conditions.

**Figure S3.**
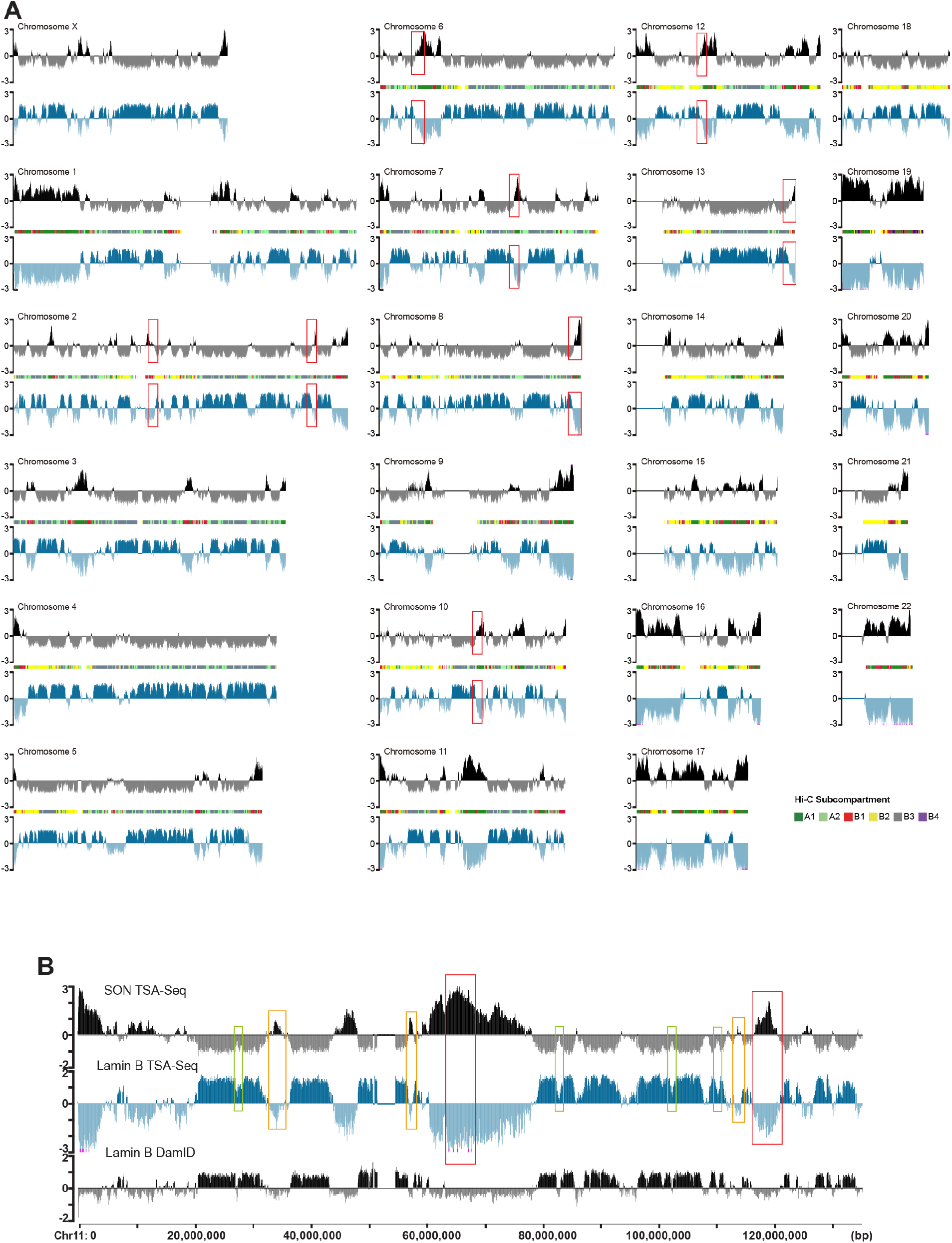
Inverse correlation between SON and lamin B TSA-Seq, as shown for all human chromosomes. (A) SON TSA-Seq (black) and lamin B TSA-Seq (blue) maps (Staining Condition 2)- y-axis shows log2 ratio of TSA normalized versus input DNA normalized reads in 20 kb bins. For comparison, Hi-C subcompartments (GM12878 cells) are shown between the SON and lamin B TSA-Seq plots (color code, bottom right). Plots are based on the hg19 assembly as visualized in the UCSC Genome Browser. Red boxes show examples of linear transitions connecting valleys (peaks) to peaks (valleys) in SON (lamin B) TSA-Seq maps. (B) SON TSA-Seq large peaks (red boxes), small peaks (orange boxes), and “peaks-within-valleys” (green boxes) correspond to Lamin TSA-Seq large valleys, shallow valleys, and dips between peaks, respectively. Top to bottom: K562 cell SON TSA-Seq, lamin B TSA-Seq, lamin B1 DamID.

**Figure S4.**
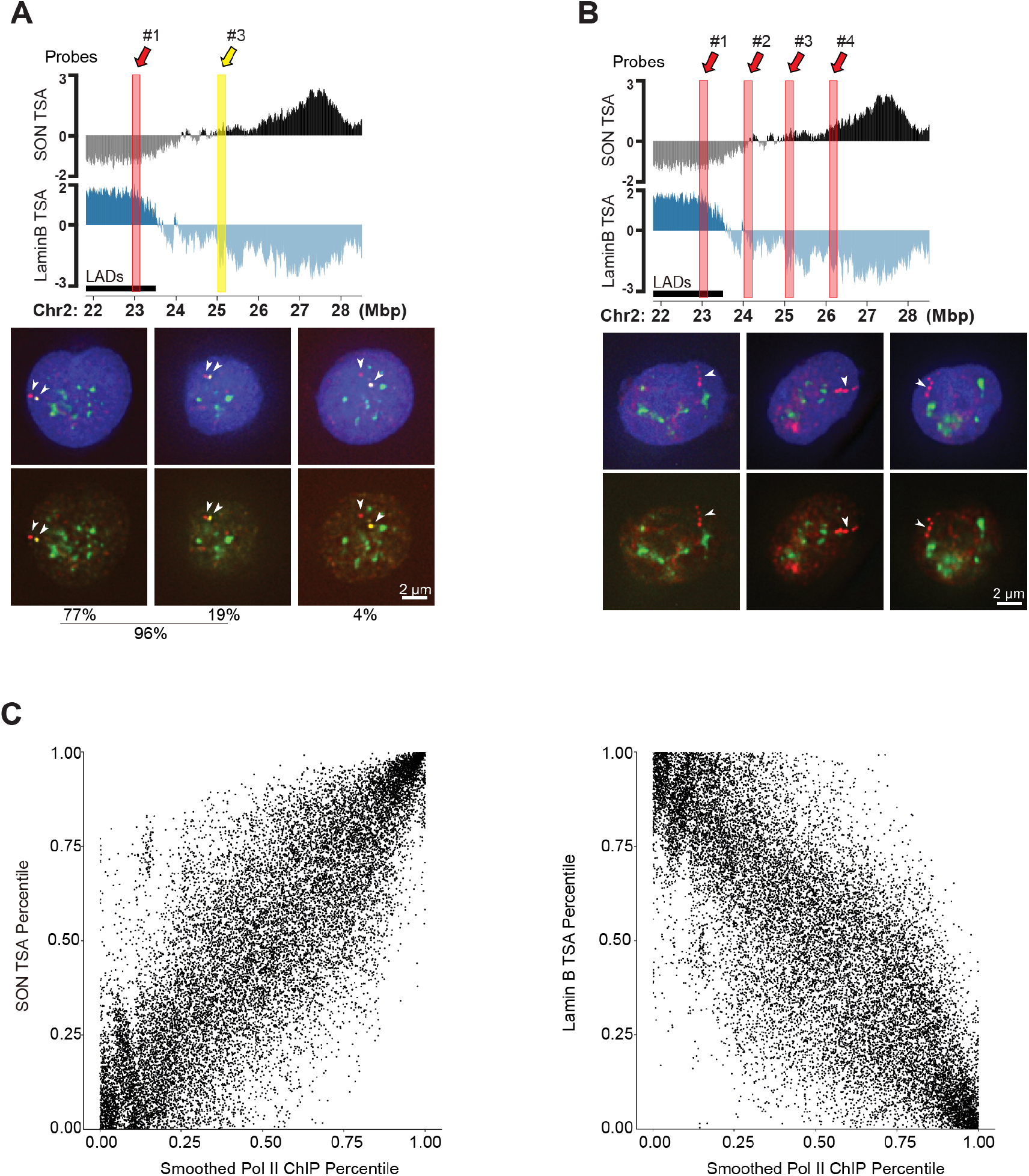
TSA-Seq maps predict both chromosome location and chromosome topology. (A-B) Top: SON and Lamin B TSA-Seq genomic plots showing locations and colors of probes used to map trajectory orientation and path. Bottom: SON speckle staining (green), probe positions (arrowheads), and DAPI (blue) DNA staining are completely (top) or partially (bottom, no DAPI) merged. Scale bars: 2 μm. (A) 77% of examples show probe 1 maps ≤0.5 μm from the periphery (in 2D) with probe 2 located more interiorly (left); 19% of examples show probe 1 closer to the nuclear periphery than probe 2 but located > 0.5 μm from the periphery (middle); 4% of examples show an indeterminate orientation (right) (N=114). (B) Chromosome trajectories point from nuclear periphery towards nuclear speckle in 69% of cases (85/124). (C) In contrast to SON versus lamin TSA-Seq (Figure 3B), Pol II ChIP signal poorly correlates with TSA-Seq. 2D scatter-plots show genome-wide relationship between SON or Lamin B TSA-Seq and pol II ChIP-Seq. Each dot represents a single 160 kb bin plotted at a position corresponding to the average TSA-Seq percentiles and 1 Mbp-smoothed normalized RNA pol II ChIP-Seq percentiles.

**Figure S5.**
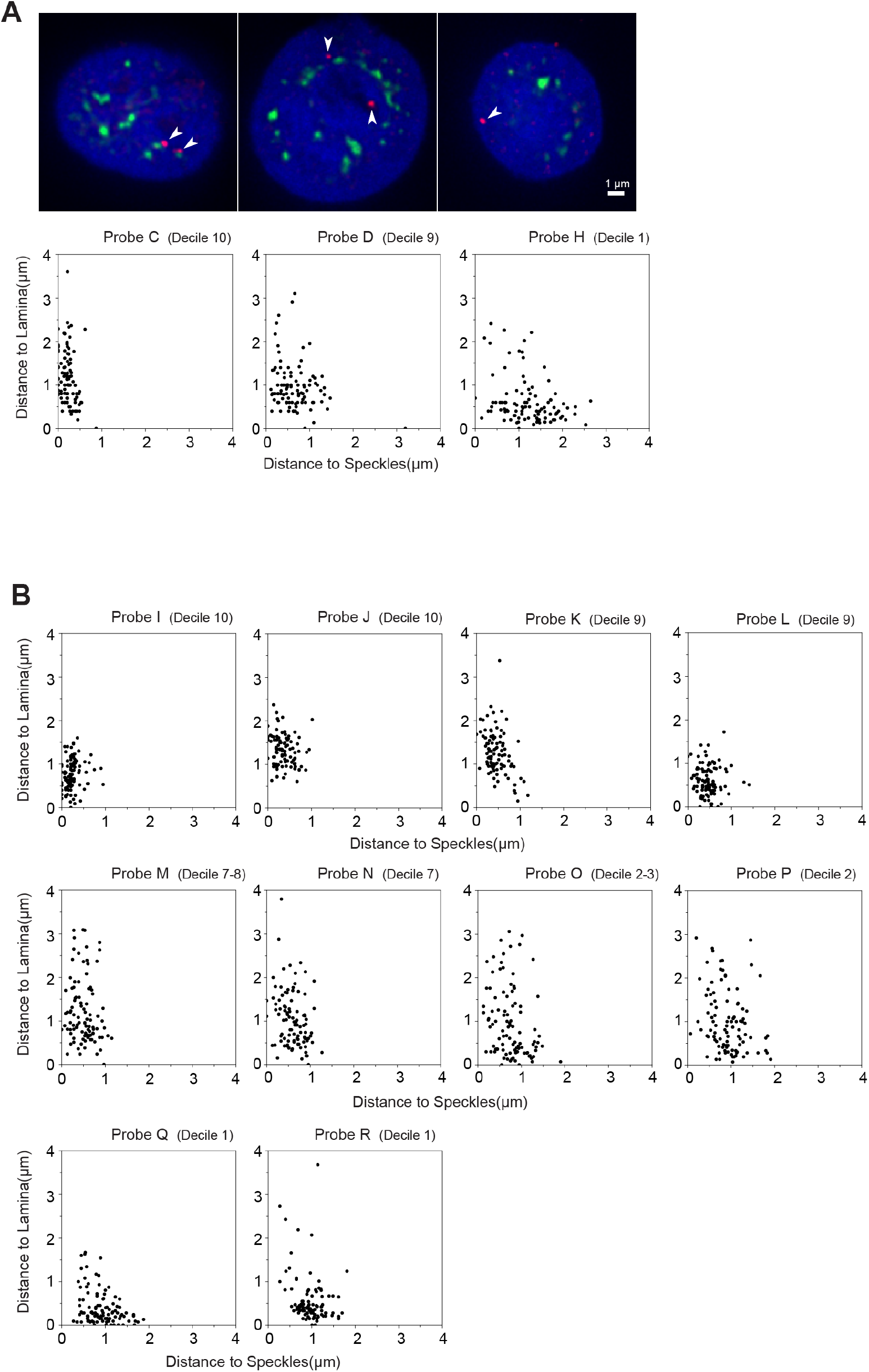
Single-cell analysis reveals that genomic regions positioned deterministically near nuclear speckles show stochastic positioning relative to the nuclear lamina. Although 2D lamin-SON TSA scatter-plots show a tight inverse correlation, 3D immuno-FISH reveals that in single cells cytological distances of a given locus from the nuclear speckle or lamina are not tightly correlated. (A) Top: DNA FISH probe location (red, arrowheads) relative to nuclear speckles (green) and nuclear DNA staining (blue, DAPI). Probes C (left, decile 10), D (middle, decile 9) and H (right, decile 1) (probes same as in Figures 2, 4, S2) are from a SON TSA-Seq peak, transition zone, and valley (overlapping with a LAD), respectively; deciles refer to SON TSA-Seq. Scale bars: 1μm. Bottom: 2D scatter-plots showing shortest distances in microns from both the nearest nuclear speckle (x-axis) and nuclear periphery (y-axis) for probes C, D, and H, measured for 100 alleles. Probe C shows a very tight distribution of distances (~0-0.5 μm) close to the nuclear speckle, but a broad distribution of distances to the periphery. Instead, Probe H shows a relatively tight distribution of distances for most alleles from the nuclear periphery (~0-0.8 μm) but a broad distribution of distances to nuclear speckles. Probe D shows distance distributions with an intermediate degree of scatter relative to both nuclear speckle and periphery. (B) 10 additional 2D scatter-plots for probes I to R (probes same as in Figures 2, 4 and S2).

**Figure S6.**
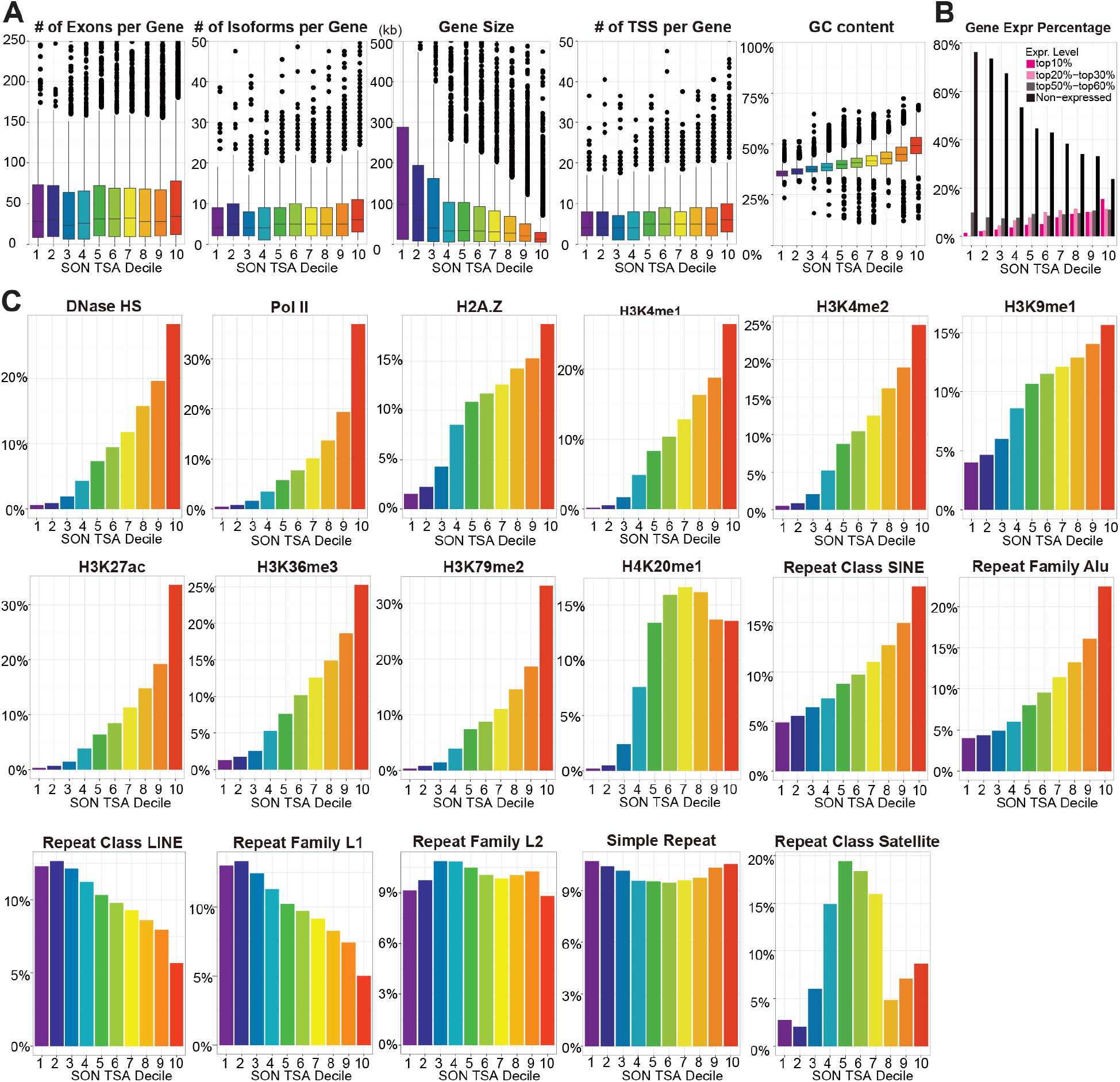
Genome features relative to SON TSA-Seq decile. (A) Box plots show genomic features in each of the SON TSA-Seq deciles. With increasing deciles, exon number and number of isoforms per gene show little change, gene size decreases, transcription start sites per gene increase slightly, and GC content increases progressively. (B) Percentage of highly expressed genes (top10%, dark pink) increases while percentage of non-expressed genes (black) decreases with increasing decile. Genes expressed at near average levels (50-60%, grey) make up a near constant fraction of expressed genes in each decile, while genes expressed in the top 20-30% percentile (light pink) are relatively depleted from the bottom 3 deciles, but otherwise show little variation. (C) Chromatin features related to transcription increase in a continuous gradient with increasing decile: DNase I hypersensitive sites peak count, RNA Pol II level, H2A.Z level, H3K4me1 peak count, H3K4me2 peak count, H3K9me1 level, H3K27ac level, H3K36me3 level and H3K79me2 peak count. A roughly binary division is seen for H4K20me1 level in the ten deciles. An increasing linear gradient relative to speckle proximity is observed for the SINE repeat class and Alu repeat family. A decreasing linear gradient is observed for the LINE repeat class and L1 repeat family. Instead, the L2 repeat families and the simple repeat class are uniformly distributed. Satellite repeats show a strong enrichment in the middle deciles. (D) Distribution of each Hi-C subcompartment type across SON TSA-Seq deciles.

**Figure S7.**
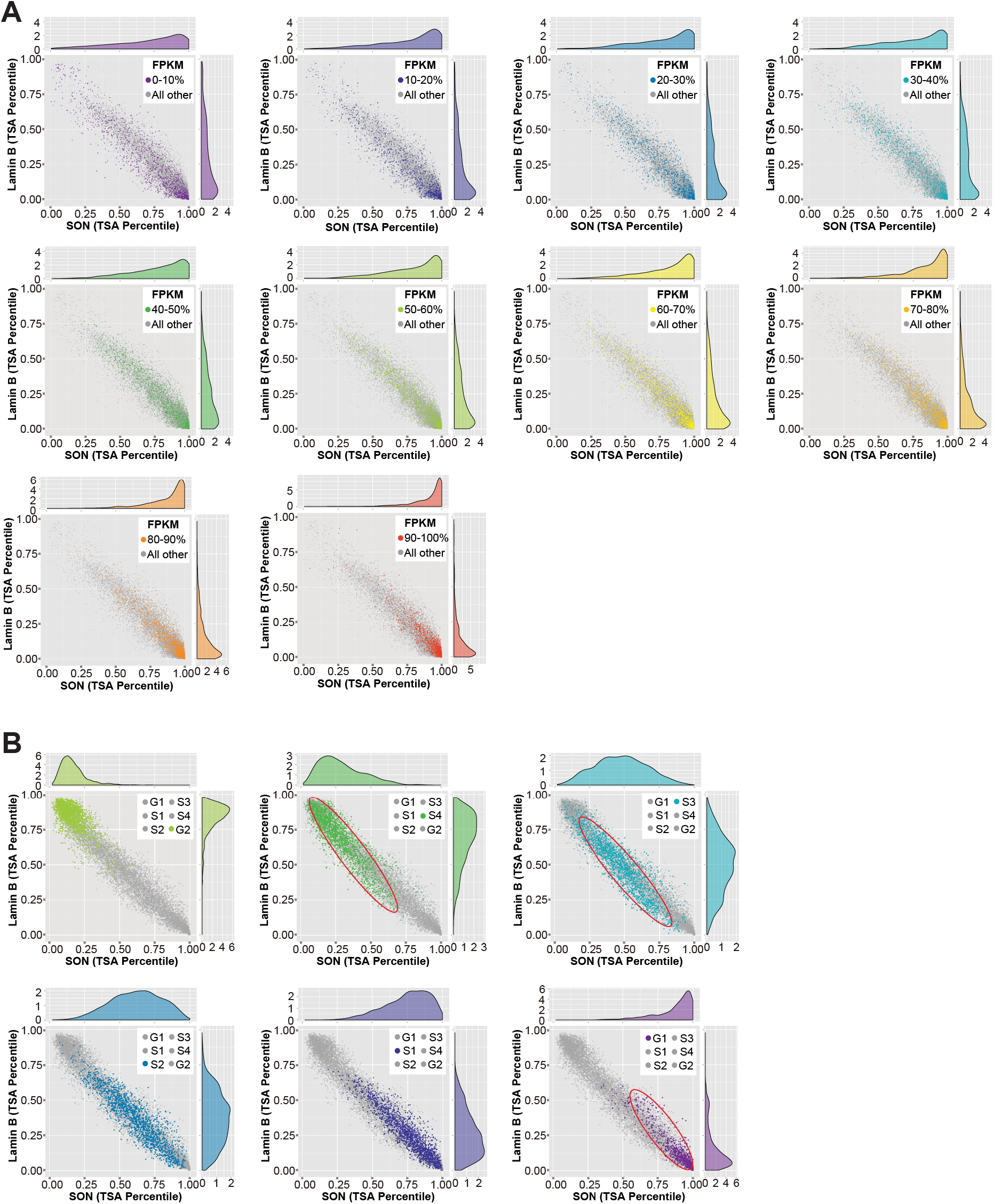
2D scatter-plots showing distribution of gene expression and DNA replication timing relative to speckle-lamin TSA-Seq axis. (A) Protein-coding expressed genes are divided into 10 deciles according to their expression level. Each gene expression decile is plotted in its own 2D lamin B versus SON TSA-Seq percentile scatter-plot, with genes in this decile appearing as colored dots; all other expressed genes appear as grey dots. Projections are shown on top and right sides. Increasing expression deciles are plotted left to right and top to bottom. As expression levels increase, gene location is skewed increasingly closer to speckles (x axis, higher SON TSA-Seq percentiles) and further from the lamina (y-axis, lower lamin B TSASeq percentiles). With the highest expression decile, there is a skewing or bias towards disproportionally higher SON TSA-Seq percentiles for regions far from the nuclear lamina and close to speckles (80-100% SON TSA-Seq percentile, 0-20% lamin B TSA-Seq percentile). (B) 6 DNA replication timing groups are plotted separately in 2D lamin B versus SON TSA-Seq percentile scatter-plots. Each dot represents a 300 kb bin. DNA replication timing shifts progressively along the speckle-lamina axis with late replicating regions concentrated in areas near the lamina but far from speckles, and early replicating regions concentrated in areas near speckles but far from the lamina. Red ellipses show the slight skewing of earlier replicating regions towards disproportionally higher SON TSA-Seq percentiles and intermediate replicating regions towards lower lamin B TSA-Seq percentiles.

## References

Agard, D.A., Hiraoka, Y., Shaw, P., and Sedat, J.W. (1989). Fluorescence Microscopy in Three Dimensions. Methods Cell Biol. 30, 353–377.

Belmont, A.S., and Bruce, K. (1994). Visualization of G1 chromosomes: A folded, twisted, supercoiled chromonema model of interphase chromatid structure. J. Cell Biol. 127, 287–302.

Bickmore W.A. (2013). The Spatial Organization of the Human Genome. Annu. Rev. Genomics Hum. Genet. 14, 67–84.

Brangwynne C.P. (2011). Soft active aggregates: mechanics, dynamics and self-assembly of liquid-like intracellular protein bodies. Soft Matter 3052–3059.

Carter, K.C., Taneja, K.L., and Lawrence, J.B. (1991). Discrete nuclear domains of poly(A) RNA and their relationship to the functional organization of the nucleus. J. Cell Biol. 115, 1191–1202.

Ching, R.W., Ahmed, K., Boutros, P.C., Penn, L.Z., and Bazett-Jones, D.P. (2013). Identifying gene locus associations with promyelocytic leukemia nuclear bodies using immuno-TRAP. J. Cell Biol. 201, 325–335.

Day, D.S., Zhang, B., Stevens, S.M., Ferrari, F., Larschan, E.N., Park, P.J., and Pu, W.T. (2016). Comprehensive analysis of promoter-proximal RNA polymerase II pausing across mammalian cell types. Genome Biol. 17, 120.

Dernburg A.F. (2011). Fragmentation and labeling of probe DNA for whole-mount FISH in Drosophila. Cold Spring Harb. Protoc. 2011, 1527–1530.

Dow, E.C., Liu, H., and Rice, A.P. (2010). T-loop phosphorylated Cdk9 localizes to nuclear speckle domains which may serve as sites of active P-TEFb function and exchange between the Brd4 and 7SK/HEXIM1 regulatory complexes. J. Cell. Physiol. 224, 84–93.

Eisenberg, E., and Levanon, E.Y. (2013). Human housekeeping genes, revisited. Trends Genet. 29, 569–574.

Fakan, S., and Puvion, E. (1980). The Ultrastructural Visualization of Nucleolar and Extranucleolar RNA Synthesis and Distribution. Int. Rev. Cytol. 65, 255–299.

Galganski, L., Urbanek, M.O., and Krzyzosiak, W.J. (2017). Nuclear speckles: molecular organization, biological function and role in disease. Nucleic Acids Res. 45, 10350–10368.

Guelen, L., Pagie, L., Brasset, E., Meuleman, W., Faza, M.B., Talhout, W., Eussen, B.H., De Klein, A., Wessels, L., De Laat, W., et al. (2008). Domain organization of human chromosomes revealed by mapping of nuclear lamina interactions. Nature 453, 948–951.

Hall, L.L., Smith, K.P., Byron, M., and Lawrence, J.B. (2006). Molecular anatomy of a speckle. Anat. Rec. Part A Discov. Mol. Cell. Evol. Biol. 288A, 664–675.

Hansen, R.S., Thomas, S., Sandstrom, R., Canfield, T.K., Thurman, R.E., Weaver, M., Dorschner, M.O., Gartler, S.M., and Stamatoyannopoulos, J.A. (2010). Sequencing newly replicated DNA reveals widespread plasticity in human replication timing. Proc. Natl. Acad. Sci. 107, 139–144.

Harrow, J., Frankish, A., Gonzalez, J.M., Tapanari, E., Diekhans, M., Kokocinski, F., Aken, B.L., Barrell, D., Zadissa, A., Searle, S., et al. (2012). GENCODE: The reference human genome annotation for the ENCODE project. Genome Res. 22, 1760–1774.

Herrmann, C.H., and Mancini, M.A. (2001). The Cdk9 and cyclin T subunits of TAK/P-TEFb localize to splicing factor-rich nuclear speckle regions. J. Cell Sci. 114, 1491–1503.

Hnisz, D., Abraham, B.J., Lee, T.I., Lau, A., Saint-André, V., Sigova, A.A., Hoke, H.A., and Young, R.A. (2013). Super-enhancers in the control of cell identity and disease. Cell 155, 934–947.

Hopman, A.H.N., Ramaekers, F.C.S., and Speel, E.J.M. (1998). Rapid synthesis of biotin-, digoxigenin-, trinitrophenyl-, and fluorochrome-labeled tyramides and their application for in situ hybridization using CARD amplification. J. Histochem. Cytochem. 46, 771–777.

Hu, Y., Kireev, I., Plutz, M., Ashourian, N., and Belmont, A.S. (2009). Large-scale chromatin structure of inducible genes: Transcription on a condensed, linear template. J. Cell Biol. 185, 87–100.

Imakaev, M., Fudenberg, G., McCord, R.P., Naumova, N., Goloborodko, A., Lajoie, B.R., Dekker, J., and Mirny, L.A. (2012). Iterative correction of Hi-C data reveals hallmarks of chromosome organization. Nat. Methods 9, 999–1003.

Ji, X., Zhou, Y., Pandit, S., Huang, J., Li, H., Lin, C.Y., Xiao, R., Burge, C.B., and Fu, X.D. (2013). SR proteins collaborate with 7SK and promoter-associated nascent RNA to release paused polymerase. Cell 153, 855–868.

Khanna, N., Hu, Y., and Belmont, A.S. (2014). HSP70 transgene directed motion to nuclear speckles facilitates heat shock activation. Curr. Biol. 24, 1138–1144.

Kim, J.H., Baddoo, M.C., Park, E.Y., Stone, J.K., Park, H., Butler, T.W., Huang, G., Yan, X., Pauli-Behn, F., Myers, R.M., et al. (2016). SON and Its Alternatively Spliced Isoforms Control MLL Complex-Mediated H3K4me3 and Transcription of Leukemia-Associated Genes. Mol. Cell 61, 859–873.

Ko, T.K., Kelly, E., and Pines, J. (2001). CrkRS: a novel conserved Cdc2-related protein kinase that colocalises with SC35 speckles. J. Cell Sci. 114, 2591–2603.

Kölbl, A.C., Weigl, D., Mulaw, M., Thormeyer, T., Bohlander, S.K., Cremer, T., and Dietzel, S. (2012). The radial nuclear positioning of genes correlates with features of megabase-sized chromatin domains. Chromosom. Res. 20, 735–752.

Landt, S.G., Marinov, G.K., Kundaje, A., Kheradpour, P., Pauli, F., Batzoglou, S., Bernstein, B.E., Bickel, P., Brown, J.B., Cayting, P., et al. (2012). ChIP-seq guidelines and practices of the ENCODE and modENCODE consortia. Genome Res. 22, 1813–1831.

Langmead, B., and Salzberg, S.L. (2012). Fast gapped-read alignment with Bowtie 2. Nat. Methods 9, 357–359.

Li H., Handsaker B., Wysoker A., Fennell T., Ruan J., Homer N., Marth G., Abecasis G., Durbin R., and 1000 Genome Project Data Processing Subgroup (2009). The Sequence Alignment/Map format and SAMtools. Bioinformatics 25, 2078–2079.

Lieberman-Aiden, E., Van Berkum, N.L., Williams, L., Imakaev, M., Ragoczy, T., Telling, A., Amit, I., Lajoie, B.R., Sabo, P.J., Dorschner, M.O., et al. (2009). Comprehensive mapping of long-range interactions reveals folding principles of the human genome. Science (80-.). 326, 289–293.

Meuleman, W., Peric-Hupkes, D., Kind, J., Beaudry, J.B., Pagie, L., Kellis, M., Reinders, M., Wessels, L., and Van Steensel, B. (2013). Constitutive nuclear lamina-genome interactions are highly conserved and associated with A/T-rich sequence. Genome Res. 23, 270–280.

Moen P.T. (2003). Repositioning of Muscle-specific Genes Relative to the Periphery of SC-35 Domains during Skeletal Myogenesis. Mol. Biol. Cell 15, 197–206.

Quinlan, A.R., and Hall, I.M. (2010). BEDTools: A flexible suite of utilities for comparing genomic features. Bioinformatics 26, 841–842.

Rao, S.S.P., Huntley, M.H., Durand, N.C., Stamenova, E.K., Bochkov, I.D., Robinson, J.T., Sanborn, A.L., Machol, I., Omer, A.D., Lander, E.S., et al. (2014). A 3D Map of the Human Genome at Kilobase Resolution Reveals Principles of Chromatin Looping. Cell 159, 1665–1680.

Schindelin, J., Arganda-Carreras, I., Frise, E., Kaynig, V., Longair, M., Pietzsch, T., Preibisch, S., Rueden, C., Saalfeld, S., Schmid, B., et al. (2012). Fiji: An open-source platform for biological-image analysis. Nat. Methods 9, 676–682.

Shopland, L.S., Johnson, C. V., Byron, M., McNeil, J., and Lawrence, J.B. (2003). Clustering of multiple specific genes and gene-rich R-bands around SC-35 domains: Evidence for local euchromatic neighborhoods. J. Cell Biol. 162, 981–990.

Sims, J.K., Houston, S.I., Magazinnik, T., and Rice, J.C. (2006). A trans-tail histone code defined by monomethylated H4 Lys-20 and H3 Lys-9 demarcates distinct regions of silent chromatin. J. Biol. Chem. 281, 12760–12766.

Solovei, I., and Cremer, M. (2010). 3D-FISH on cultured cells combined with immunostaining. Methods Mol. Biol. 659, 117–126.

Spector, D.L., and Lamond, A.I. (2011). Nuclear speckles. Cold Spring Harb. Perspect. Biol. 3, a000646.

van Steensel, B., and Belmont, A.S. (2017). Lamina-Associated Domains: Links with Chromosome Architecture, Heterochromatin, and Gene Repression. Cell 169, 780–791.

Takizawa, T., Meaburn, K.J., and Misteli, T. (2008). The meaning of gene positioning. Cell 135, 9–13.

Vogel, M.J., Peric-Hupkes, D., and van Steensel, B. (2007). Detection of in vivo protein–DNA interactions using DamID in mammalian cells. Nat. Protoc. 2, 1467–1478.

Wang, G., Achim, C.L., Hamilton, R.L., Wiley, C.A., and Soontornniyomkij, V. (1999). Tyramide Signal Amplification Method in Multiple-Label Immunofluorescence Confocal Microscopy. Methods 18, 459–464.

Wang, Z., Zang, C., Rosenfeld, J.A., Schones, D.E., Barski, A., Cuddapah, S., Cui, K., Roh, T.-Y., Peng, W., Zhang, M.Q., et al. (2008). Combinatorial patterns of histone acetylations and methylations in the human genome. Nat. Genet. 40, 897–903.

Wartlick, O., Kicheva, A., and González-Gaitán, M. (2009). Morphogen gradient formation. Cold Spring Harb. Perspect. Biol. 1, a001255.

Xing, Y., Johnson, C. V., Moen, P.T., McNeil, J.A., and Lawrence, J.B. (1995). Nonrandom gene organization: Structural arrangements of specific pre-mRNA transcription and splicing with SC-35 domains. J. Cell Biol. 131, 1635–1647.

Zhu, L., and Brangwynne, C.P. (2015). Nuclear bodies: the emerging biophysics of nucleoplasmic phases. Curr. Opin. Cell Biol. 34, 23–30.

